# Higher-Throughput Proteome Profiling Enabled by Parallelized Pre-Accumulation and Optimized Ion Processing in the Orbitrap Astral Zoom Mass Spectrometer

**DOI:** 10.1101/2025.07.14.664789

**Authors:** Ulises H. Guzman, Martin Rykar, Ivo A. Hendriks, Hamish Stewart, Eduard Denisov, Bernd Hagedorn, Johannes Petzoldt, Arne Kreutzmann, Yannick Mueller, Tabiwang N. Arrey, Immo Colonius, Ole Østergaard, Claire Koenig, Julia Kraegenbring, Kyle L. Fort, Erik Couzijn, Jan-Peter Hauschild, Daniel Hermanson, Vlad Zabrouskov, Christian Hock, Eugen Damoc, Jesper. V. Olsen

## Abstract

High-throughput proteomics is critical for understanding biological processes, enabling large-scale studies such as biomarker discovery and systems biology. However, current mass spectrometry technologies face limitations in speed, sensitivity, and scalability for analyzing large sample cohorts. The Thermo Scientific™ Orbitrap™ Astral™ Zoom mass spectrometer (MS) was developed to address these limitations by improving acquisition speed, ion utilization, and spectral processing, which are all essential for advancing proteome depth in high-throughput proteomics. The Orbitrap Astral Zoom MS achieves ultra-fast MS/MS scan rates of up to 270 Hz with enhanced ion utilization through pre-accumulation, enabling the identification of ∼100,000 unique peptides and >8,400 proteins in a single 300 samples-per-day (SPD) analysis of human cell lysate. The optimized system reduces analysis time by 40%, achieves near-complete proteome coverage (>12,000 proteins) in 2.7 hours, and enables ultra-high-throughput workflows, identifying >7,000 proteins in a 500 SPD method with exceptional reproducibility (Pairwise Pearson correlations >0.99). These advancements establish the Orbitrap Astral Zoom MS as a new benchmark in proteomics, significantly enhancing speed, sensitivity, and scalability, paving the way for routine large-scale proteome studies with applications in clinical research and systems biology.

**Teaser:** High-Speed Human Proteome Analysis using the Orbitrap Astral Zoom Mass Spectrometer

## Introduction

Biological systems are complex networks essential for organismal function, ranging from individual cells to complex ecosystems. They are inherently dynamic and their rapid response to perturbations are influenced by a variety of factors, such as genetic background and environmental conditions^1,2^. To understand their behavior, it is essential to systematically assess the proteome states across diverse populations, disease contexts, and model systems under varying conditions^3,4^. Proteomics based on peptide sequencing by online liquid chromatography tandem mass spectrometry (LC-MS/MS) is generally the method of choice for such applications. Recent technological advancements in MS-based proteomics technologies have facilitated remarkable proteome depth, sensitivity, and throughput^5–8^. However, systems biology studies, biomarker discovery from large clinical research cohorts, or drug library screenings by MS-based proteomics, are dependent on deep proteome coverage with high reproducibility across hundreds or thousands of samples. Therefore, further scaling of such proteomics experiments is crucial, not only to decrease MS measurement time but also to generate high-quality, comprehensive datasets suitable for advanced statistical analyses and modelling, including machine learning^9,10^. Traditionally, comprehensive MS projects require days, weeks, or even months of LC-MS instrument time. Addressing these demands requires the shortening of chromatographic gradients, while preserving both sensitivity and proteome coverage. As a result, mass spectrometers must operate at higher acquisition rates to process the same number of analytes within reduced time frames^11^, while ensuring sufficient selectivity to resolve co-eluting peptides and preserve quantitative performance. State-of-the-art proteomics-grade MS systems operated in data-independent acquisition (DIA) mode are capable of acquiring more than one hundred high-resolution accurate mass DIA-MS/MS scans per second, which enabled the development of fast acquisition methods^12^. For instance, the Thermo Scientific™ Orbitrap™ Astral™ MS can achieve acquisition rates of ∼200 Hz^13–15^, and narrow (2 Th) isolation windows that have proven effective for DIA applications, despite reduced utilization of the ion beam^8^. Although these capabilities are significant, further improvements in speed and sensitivity can be achieved by optimizing the ion processing stage and further parallelizing ion handling during the instrument’s overheads. The operation of the Orbitrap Astral MS is highly parallelized, with multiple ion packets processed simultaneously to decouple ion accumulation time from downstream processing stages. Nevertheless, it still suffers from almost 2 ms scan-to-scan overhead, which substantially impacts duty cycle at maximum 200 Hz operational scanning speed equivalent of ∼5ms acquisition time per MS/MS spectrum^13^. Here, we present an optimized ion processing stage and enhanced spectral processing algorithm^16^ on an Orbitrap Astral Zoom MS to improve the acquisition rate, duty cycle, and spectral processing for deep proteome profiling. The optimized ion processing method, in conjunction with narrow-window DIA (nDIA) and the enhanced spectral processing algorithm, enables ultra-fast MS/MS scans rates of up to 270 Hz, preserving the high selectivity of narrow DDA-like isolation windows without sacrificing ion accumulation time and spectral quality. Compared to the Orbitrap Astral MS^17^, the Orbitrap Astral Zoom MS identified more peptides and proteins in half the LC gradient time with ∼100,000 unique peptides and ∼8400 protein-coding genes (hereafter referred to as protein groups or PGs) with a 300 SPD method and >7000 unique proteins from a 500 SPD LC-MS/MS method. Similar improvements in analytical depth were observed with extended gradients, such as 24 SPD, enabling the identification of >10,200 protein groups and >200,000 unique peptides. Furthermore, these optimizations accelerated the acquisition of near-complete proteomes (> 12,000 PGs), reducing the analysis time for 34 high-pH fractions by 40%. This advancement allowed for the acquisition of a full proteome in just 2.7 hours. This approach also provides sufficient depth to confidently identify and localize more than 2000 phosphosites without enrichment, as well as ∼ 1,600 splicing variants.

## Results

### Optimized ion processing stage enables increased ion beam utilization and faster MS/MS acquisition

The Orbitrap Astral mass spectrometer pairs a quadrupole with both a Thermo Scientific^TM^ Orbitrap^TM^ analyzer and a Thermo Scientific^TM^ asymetric track lossless (Astral) analyzer providing >200Hz MS/MS scanning speed, high resolving power and sensitivity, and low-ppm mass accuracy. This configuration enables the sensitivity and acquisition speed to routinely perform DIA analyses with quadrupole isolation window width similar to data-dependent acquisition (DDA), blurring the contrast between DIA and DDA^17^. Here, we introduce the Orbitrap Astral Zoom MS hardware and software modifications that improve MS/MS scan speed, increase ion beam utilization and sensitivity, and improve the MS/MS peak deconvolution across the full m/z range. (Fig. 1A). Briefly, faster ion filter quadrupoles (blue) have been introduced to enable higher acquisition rates. To recover duty cycle lost due to overheads during ion transfers, enhance the ion beam utilization and increase sensitivity, a parallel ion pre-accumulation stage (Pre-Ac) has been added in the bent trap (pink) upstream of the mass selection quadrupole. To reduce timing delay overheads, stronger ion shuttling axial fields in the ion routing multipole (IRM) and more stable control hardware were implemented. Moreover, faster ion transfer in the ion processor (yellow) has been implemented to increase acquisition rates, and the introduction of a relay prism in the Astral analyzer (purple) facilitates higher-resolution scans. Collectively, these modifications enable an optimized ion processing throughout the Orbitrap Astral Zoom compared to the Orbitrap Astral MS (Supplementary Fig. 1A); from the inject flatapole, through the bent trap, mass filter quadrupole, ion routing multipole (IRM), ion processor, and Astral analyzer. This reduces the median scan-to-scan time from 4.32 ms in the Orbitrap Astral to 3.49 ms in the Orbitrap Astral Zoom, improving overall acquisition speed (Fig. 1B), enabling 270 Hz acquisition rates. Moreover, the Pre-Ac stage occurring in the bent trap (a curved quadrupolar PCB-mounted ion guide with a superimposed DC gradient generated by a PCB printed electrode series^18^), operates concurrently while downstream devices process the previous ion packet in a on-demand automatic gain controlled (AGC) fashion (Fig.1C, Supplementary Fig. 1B). This approach adds approximately 1 ms of ion accumulation without increasing the overall cycle time. This enhances duty cycle and sensitivity at constant scan speed, sustains sensitivity at higher MS/MS acquisition rates. In addition, an enhanced spectral processing (ESP) algorithm was implemented across the whole mass range to deconvolute overlapping spectral features in MS/MS scans to better resolve feature-rich spectra, rather than rely entirely on the analyzer’s high resolving power to baseline separate different m/z peaks (Supplementary Fig. 1C). Briefly, this method segments the ion signal trace based on arrival times, applying adaptive filters tuned to expected ion arrival distributions to each segment, identifying ion peaks in the filtered data, and determining their characteristics—thereby reducing misinterpretation of multimodal peaks as overlapping ions^16^. Enabling this feature on the Orbitrap Astral Zoom MS resulted in a median 4-fold increase in the number of MS/MS spectral peaks compared to the Orbitrap Astral MS at 300 SPD gradients (Fig. 1D). Finally, to extend the dynamic range of Orbitrap MS1 scans, an enhanced Dynamic Range (eDR) mode acquisition strategy was implemented. This feature overcomes the constraints imposed by intense ion populations in filling the ion routing multipole and transversing the C-trap, which typically hinder detection of low-abundance analytes by multiplexing injections with variable ion accumulation times across different m/z ranges (Supplementary Fig. 1D). This approach selectively attenuates high-intensity m/z regions while amplifying low-intensity species. The eDR mode segments the MS1 survey scan into predefined regions with tailored ion accumulation targets, substantially increasing the number of detected peaks and improving signal-to-noise ratios for trace analytes. Using the eDR mode on the Orbitrap Astral Zoom MS resulted in up to a 20-fold improvement in signal-to-noise compared to the standard MS1 full scan and increased the median number of observed peaks in MS1 full scan by over 20 % from 3742 to 4571 (Fig. 1E).

**Fig. 1.**
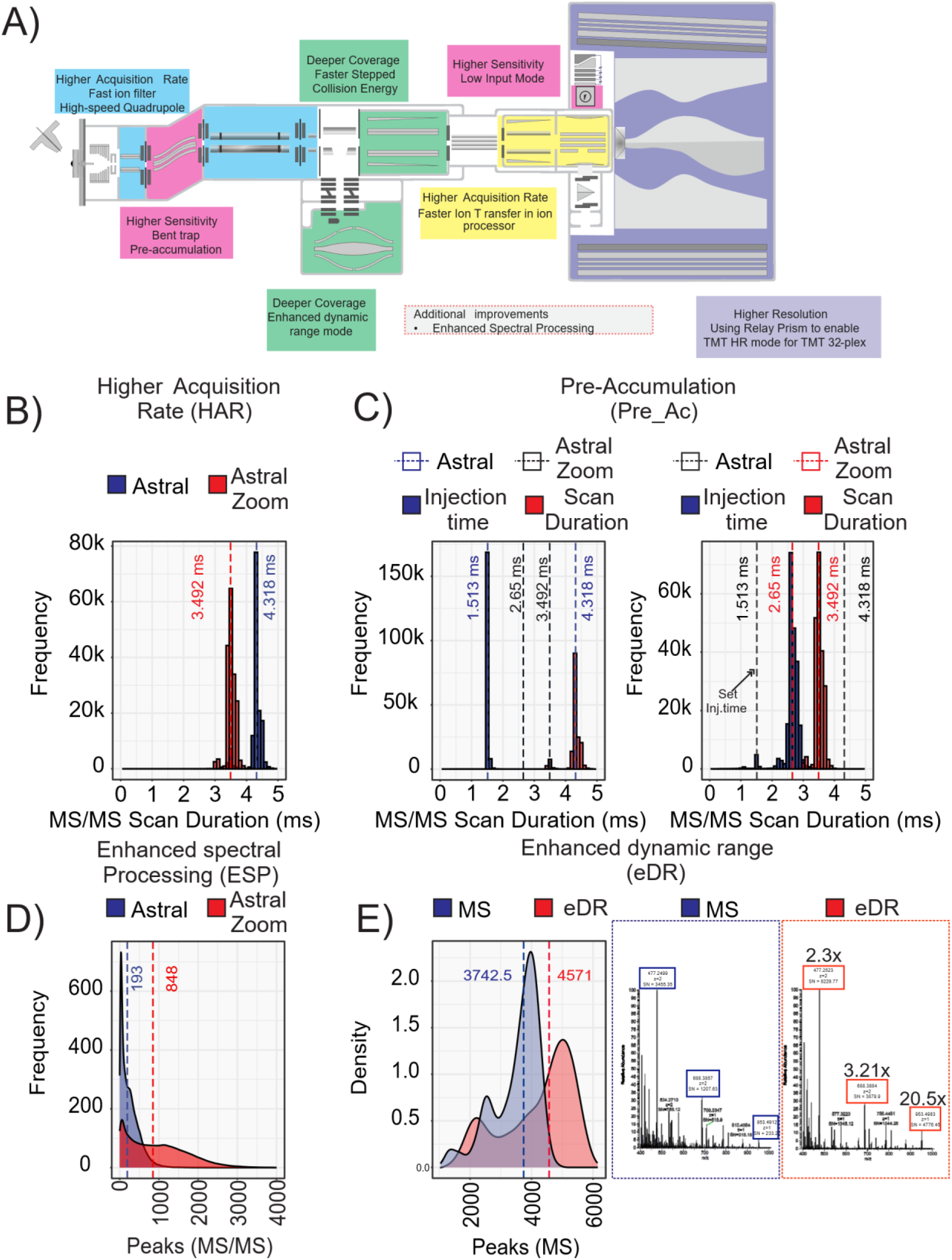
Schematic of the Orbitrap Astral Zoom mass spectrometer and enhanced ion processing features. **A)** Hardware overview of the Orbitrap Astral Zoom (OAZ) instrument, highlighting new features over the Orbitrap Astral MS (OA). Median values are indicated by dashed lines. **B)** MS/MS scan duration comparison between OA (blue)and OAZ (red) with high acquisition rate (HAR) enabled in OAZ **C)** Effective injection time comparison between OA and OAZ with the pre-accumulation feature (Pre-Ac) implemented in OAZ. **D)** Density plot showing the total number of MS/MS peaks detected in OA and OAZ with the enhanced spectral processing (ESP) algorithm enabled.

### Benchmarking the Impact of Hardware and Software Enhancements with the Orbitrap Astral Zoom MS

To determine the optimal ion injection time for maximizing scan speed and ion beam utilization, we performed a titration experiment in which the injection time in the ion routing multipole was incrementally increased from 1.0 to 5.5 ms in 0.1 ms steps, while operating the MS instrument in narrow-window DIA mode with 2 Th quadrupole isolation across a mass range of m/z 380-980 and 200 ng of HEK293 tryptic digest was analyzed using a high-throughput 300 SPD method. We evaluated the DIA acquisition speed [Hz] as a function of injection time on the Orbitrap Astral Zoom MS and compared it to the Orbitrap Astral MS (Figure 2A). This analysis showed that the fastest scan rates of ∼270 Hz can be achieved on the Orbitrap Astral Zoom MS when injection time is restricted to 1.5 ms or less, whereas the Orbitrap Astral MS can reach scan rates of ∼220 Hz when restricting the IT to 2.5 ms or less. Notably, when adding the 1 ms free bent trap pre-accumulation to the optimal 1.5 ms IT on the Orbitrap Astral Zoom MS, the resulting effective ion accumulation time matches perfectly to the optimal 2.5 ms IT on the Orbitrap Astral MS. As expected, the Orbitrap Astral Zoom MS outperformed the Orbitrap Astral MS instrument at every tested injection time, delivering at least a 10% gain in the number of peptides across all conditions.

**Fig. 2.**
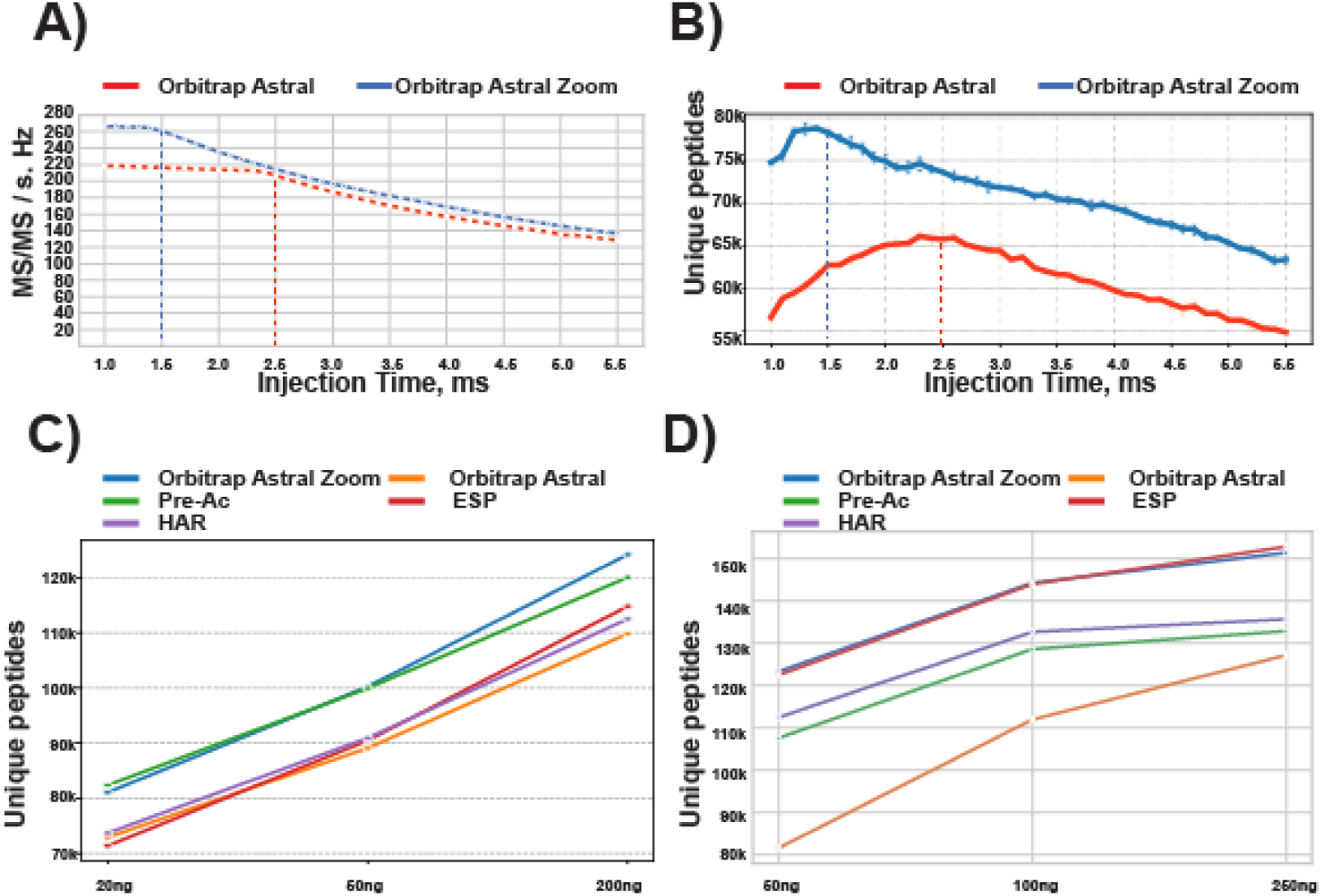
Individual benchmarking of performance-enhancing features in the Orbitrap Astral Zoom mass spectrometer. **A)** MS/MS scan rate as a function of ion injection time for the Orbitrap Astral Zoom MS and Orbitrap Astral MS. Optimal injection times for each system are indicated by dashed lines. **B)** Number of unique peptides identified as a function of ion injection time for the Orbitrap Astral Zoom MS and Orbitrap Astral MS, with optimal values indicated by dashed lines. **C)** Contribution of individual hardware and software features to peptide identification yield using fast LC gradients (90 SPD), with injection time restricted to 1.5 ms and 2 Th DIA isolation windows. Shown are the Orbitrap Astral Zoom MS (blue), Orbitrap Astral MS (orange), pre-accumulation (Pre-Ac; green), enhanced spectral processing (ESP; red), and high acquisition rate mode (HAR; violet). **D)** Same analysis as in panel C, performed using longer LC gradients (50 SPD), with injection time restricted to 6 ms and 4 Th DIA isolation windows.

Furthermore, the fastest scanning methods also yielded the highest identification rates, with the Orbitrap Astral Zoom MS identifying ∼80,000 unique peptides compared to ∼67,000 unique peptides identified on the Orbitrap Astral MS, corresponding to a ∼20% increase in peptides when optimally operating the Orbitrap Astral Zoom MS (Fig. 2B). To systematically evaluate the individual contributions of the new hardware and software features implemented in the Orbitrap Astral Zoom MS, we independently benchmarked each feature and compared the results to Orbitrap Astral and Orbitrap Astral Zoom MS. Briefly, we performed a series of DIA experiments in which each feature was individually enabled under two acquisition strategies: a high-sensitivity method (6 ms injection time, 4 Th isolation windows, 50 SPD) and a fast-scanning method (1.5 ms injection time, 2 Th windows, 90 SPD), using a range of peptide input amounts (20, 50, and 200 ng for the fast-scanning method; 50, 100, and 250 ng for the high-sensitivity method). The features assessed included pre-accumulation (Pre-Ac), higher acquisition rate (HAR), and enhanced spectral processing (ESP). Briefly, in the fast-scanning method, pre-accumulation emerged as the primary contributor to the performance gains observed for the Orbitrap Astral Zoom MS (Fig. 2C, Supplementary Fig. 2 A,B). This was reflected in both the total number of unique peptides identified and the number of peptides quantified with a coefficient of variation (CV) <20%, with Pre-Ac alone accounting for the majority of the increase and achieving performance levels nearly equivalent to the Orbitrap Astral Zoom MS with all features enabled. This can be explained by the fact that fast-scanning methods restrict the maximum injection time to 1.5 ms, thereby limiting the number of ions accumulated prior to analysis in the Astral analyser^19^. The resulting reduction in ion population can negatively affect spectral quality and sensitivity, particularly for low-abundance peptides. Consequently, the Pre-Ac feature yields a higher number of peptide identifications at lower input amounts compared to Orbitrap Astral MS and other individual features, as at shorter injection times, Pre-Ac enabled a relative increase in ion accumulation time of approximately 66%, resulting in higher average scan intensities than those achieved by enabling either HAR or ESP alone (Supplementary Fig. 2E). Meanwhile, enabling HAR and ESP individually provides modest improvements in unique peptide identifications compared to the Orbitrap Astral MS, particularly at mid-to-high peptide loads, highlighting the synergistic advantage of combining all features in the Orbitrap Astral Zoom MS. Conversely, when operating the instrument with a high-sensitivity method, the ESP algorithm accounted for the majority of the observed performance gains (Fig. 2D, Supplementary Fig. 2 C,D). The primary difference in the contributing factors to MS performance between the fast-scanning and high-sensitivity methods likely reflects two key factors. First, the longer injection time allowed in the high-sensitivity method reduces the relative contribution of scan overheads to the total MS/MS scan-to-scan time duration. This improves the duty cycle efficiency and ion utilization, reducing the contribution of the pre-accumulation feature which leverages overhead time. Second, the increased transmission resulting from a wider DIA isolation window and longer injection times facilitates the acquisition of richer-feature chimeric MS/MS spectra. These spectra can be more effectively resolved when the ESP algorithm is enabled, rather than solely relying on the high resolving power of the Astral analyzer. Consequently, these results in search engines being able to match a greater number of fragments per peptide (Supplementary Fig. 2F). Furthermore, compared to the Orbitrap Astral MS, the advancements in both hardware and software within the Orbitrap Astral Zoom MS have resulted in a 51% increase in the number of unique peptides identified when analyzing a 50 ng HEK293 tryptic digest. The contributions of individual features to this improvement include pre-accumulation (31%), high acquisition rate (38%), and enhanced spectral processing (49%), relative to the performance of the Orbitrap Astral MS (Supplementary Fig. 2G). These results illustrate the synergistic effects of all features available with the Orbitrap Astral Zoom MS even when the instrument is operated under high-sensitivity methods.

### Evaluation of the Accuracy and Precision of LFQ on the Orbitrap Astral Zoom MS under Rapid Scanning rates

To evaluate the quantitative performance of the Orbitrap Astral Zoom MS compared to the Orbitrap Astral MS in terms of precision and accuracy under high-speed scanning conditions (270 and 220 Hz respectively), we employed a label-free three-species proteome mixture, for which we mixed yeast and E.coli proteins in different ratios while keeping human proteins constant (Fig. 3A). The samples were acquired using a 50 SPD method with 2 Th isolation windows, and the maximum injection time was capped at 1.5 ms for both instruments. Notably, the Orbitrap Astral Zoom MS consistently identified an average ∼14,000 PG, exceeding the Orbitrap Astral MS by >1,000 PG (Fig. 3B). More importantly, the quantitative accuracy of the Orbitrap Astral Zoom MS proved to be on par with or even exceeded the quantitative accuracy of Orbitrap Astral MS based on the absolute median standard deviations of protein ratios compared to the theoretical ones (Fig.3C). This analysis suggested that the faster scanning methods, which yield a higher number of PG identifications, do not have a negative impact on the quantitative performance. Moreover, when the maximum injection time for the Orbitrap Astral Zoom MS was limited to 2.5 ms— the optimal injection time established for the Orbitrap Astral MS (Supplementary Fig. 3A)—the Orbitrap Astral Zoom MS exhibited superior performance compared to the Orbitrap Astral MS at the same injection time. This advantage was also observed when the Orbitrap Astral capped to 2.5 ms is compared to The Orbitrap Astral Zoom with the injection time restricted to 1.5 ms, leading to enhanced proteome coverage and quantitation. This improvement can be attributed to the higher scanning speed of the Orbitrap Astral Zoom MS, which allows for the detection and utilization of more peptides for quantitation than the Orbitrap Astral MS (Supplementary Fig. 3B). Furthermore, the increased scan speed of the Orbitrap Astral Zoom MS contributed to greater data completeness across all conditions (Supplementary Fig. 3C).

**Fig. 3.**
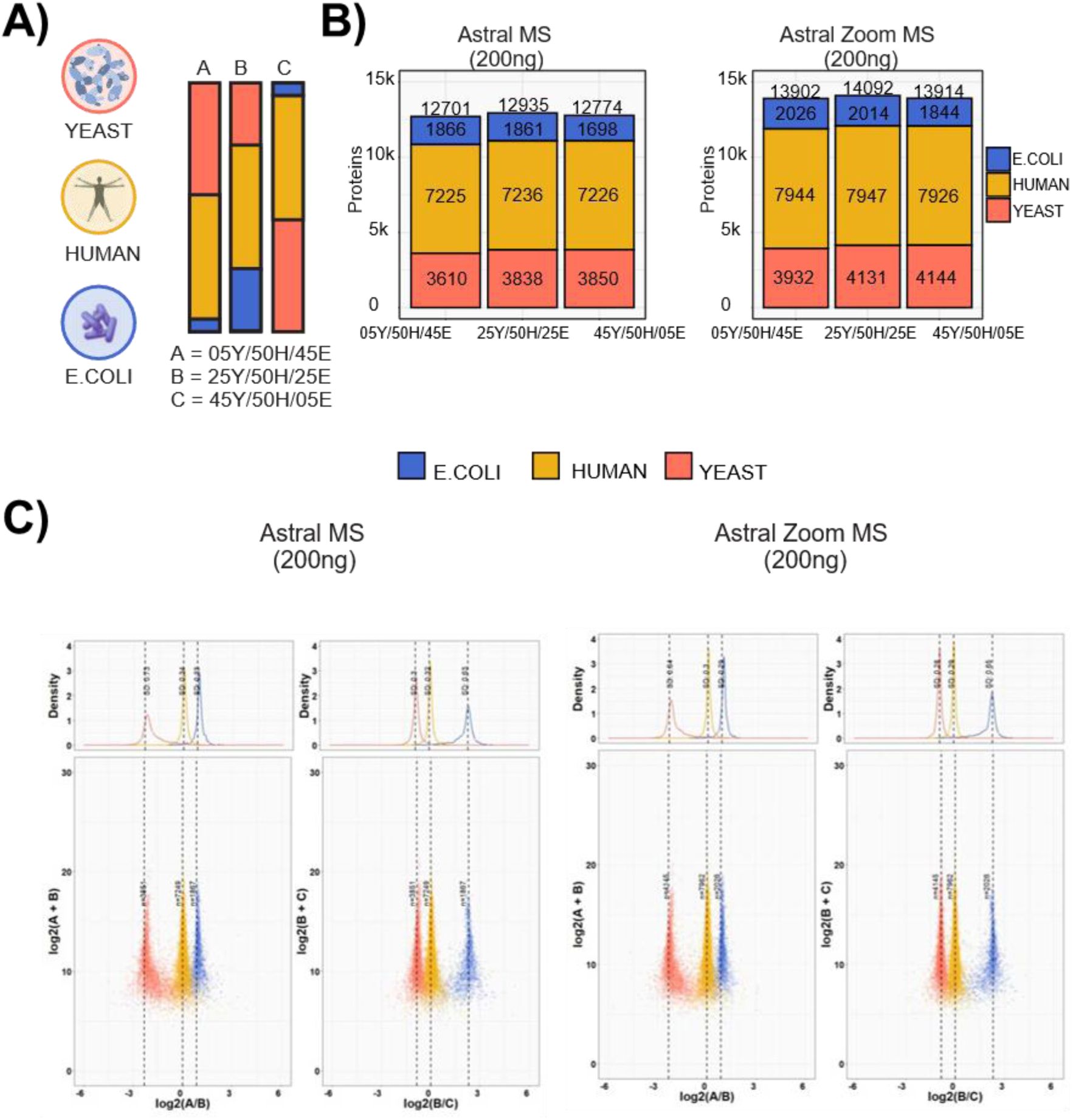
Quantitative Assessment of LFQ Accuracy and Precision on the Orbitrap Astral Zoom MS under High-Speed Scanning Regimes. **A)** Graphical representation of the experimental design. Tryptic peptides from three-species were combined in three distinct ratios (E05H50Y45, E25H50Y25 and E45H50Y05). Samples were processed using the Orbitrap Astral MS and Orbitrap Astral Zoom MS in technical triplicates, employing a 1.5-ms maxIT and 2-Th window size method. The loading amounts were 200 ng. **B)** Number of proteins identified from the three species in each sample. For the Orbitrap Astral MS and Orbitrap Astral Zoom MS. **C)** log- transformed ratios of quantified proteins. Scatter plots for all runs over the log- transformed protein intensities are displayed at the bottom, while density plots are on the top. Colored dashed lines represent expected log2(A/B) values for proteins from humans (yellow), yeast (orange) and E. coli (blue). Standard deviations are displayed on the density plots.

### Enhancing MS1 Dynamic Range via Multiplexed Ion Injections with Differential accumulation time across m/z segments

Although DIA has become the method of choice for single-shot proteomics, DDA remains valuable for specific applications including the generation of spectral libraries^20^, tandem mass tag (TMT)-based experiments^21–23^ and inmunopeptidomics^24^, among others. However, DDA is inherently constrained by the number of precursors that can be selected for fragmentation in each cycle. This selection is typically biased towards the most intense ions species observable in the full MS survey scan, which can lead to the suppression of lower-abundance peptide species due to dynamic range limitations of the C-trap^25^. When ion signals are sufficiently suppressed, they may fall below the threshold for MS/MS triggering, resulting in missed peptide identifications and reduced proteome coverage. To overcome this limitation and expand both ion sampling and dynamic range in full-scan MS, we implemented the enhanced dynamic range (eDR) mode on the Orbitrap Astral Zoom MS. This approach partitions the MS1 mass range into predefined wide SIM windows, each acquired with differential ion injection times, and merges them into a single Orbitrap full scan, while MS/MS spectra are acquired in parallel using the Astral analyzer. To evaluate the performance of this mode, we performed a series of DDA experiments with a 100 SPD method benchmarking the Orbitrap Astral Zoom MS against Orbitrap Astral Zoom with eDR mode enabled and the Orbitrap Astral MS. As expected, although the Orbitrap Astral Zoom MS incorporates features such as pre-accumulation and higher acquisition rates that improve ion utilization and scanning efficiency (data dependent MS/MS scanning speed up to 200 Hz, Supplementary Fig. 4B), these enhancements cannot fully overcome DDA’s inherent limitation. As a result, only modest improvements in peptide and protein group identification rates were observed relative to the Orbitrap Astral MS. However, enabling the eDR mode on the Orbitrap Astral Zoom MS resulted in a substantial increase in precursor identifications, yielding ∼15,000 and ∼30,000 additional precursors compared to standard and wide-window DDA methods, respectively, on the Orbitrap Astral MS (Fig. 4A). This improvement was a consequence of eDR MS1 consistently triggering ∼150 MS/MS scans per second, yielding > 80k MS/MS spectra across the peptide elution window at a fixed MS2 injection time of 3 ms. By contrast, conventional, DDA methods failed to maintain comparably high scan rates throughout the gradient (Figure 4B, Supplementary Fig. A,C).

**Fig. 4.**
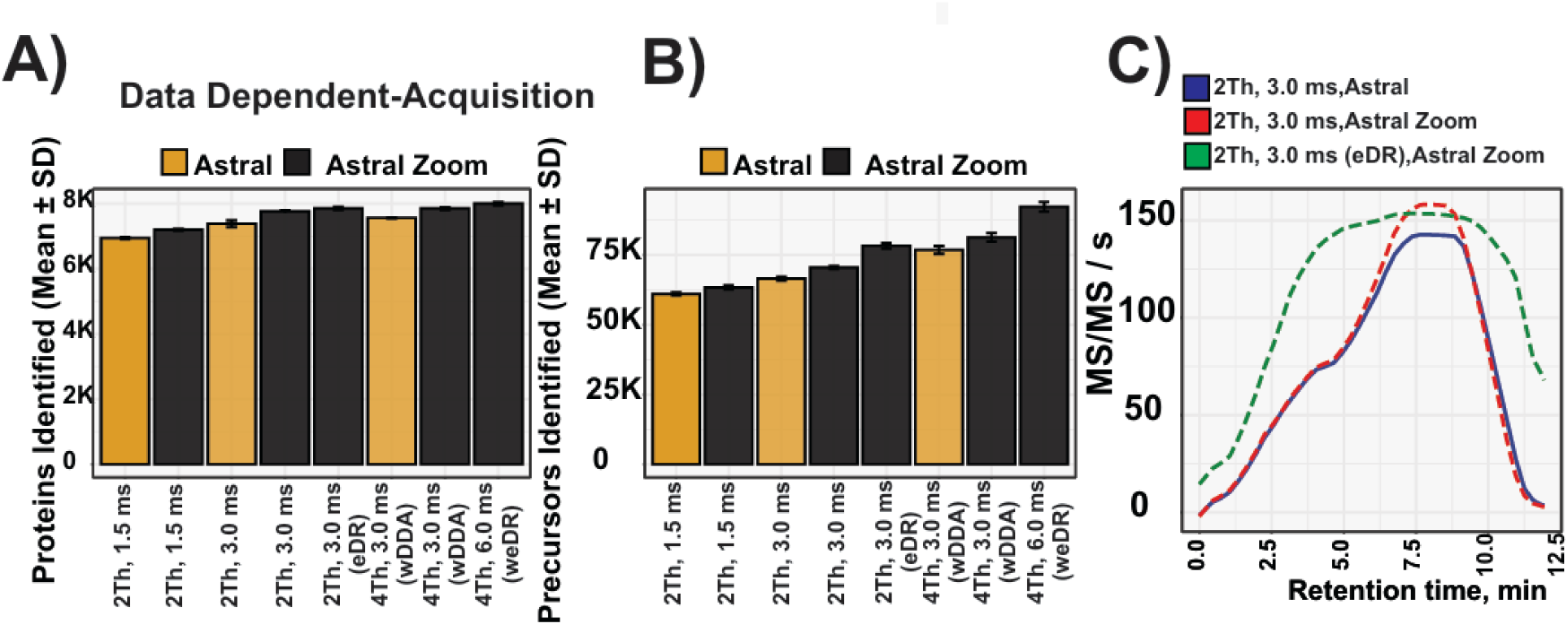
Benchmarking enhanced MS1 Dynamic Range in DDA Mode. **A)** Data-dependent acquisition (DDA) performance at the protein group level comparing the Orbitrap Astral Zoom MS and the Orbitrap Astral MS at 100 samples per day (SPD) using 10 ng HEK293 tryptic peptides across four MS methods, with and without using enhanced dynamic range (eDR) mode. **B)** Corresponding analysis at the peptide level. **C)** MS/MS scan rate (scans per second) over the LC gradient. (Data were processed using DIA-NN v1.9.)

### Optimized Fractionation Strategy for Rapid and Comprehensive Human Proteome Profiling via Multi-Shot proteomics

Based on next-generation sequencing data, human cancer cell lines like HeLa and HEK293 express ∼12,500 protein-encoding genes. We have previously demonstrated that using deep peptide fractionation via offline high-pH (HpH) reversed-phase chromatography in combination with online low-pH LC-MS/MS, it is possible to identify essentially all of the expressed proteins in a cell line^26^. In the first implementation of this strategy, it took 34.5 h of MS measurement time to achieve near-complete proteome coverage, but with the Orbitrap Astral MS this was reduced to just ∼4.5 h by analyzing 34 offline HpH fractions of a tryptic digest of HEK293 cells with a 180 SPD nDIA^17^. This represented a 7x improvement in throughput, enabling deep proteome profiling with dramatically increased efficiency. To test whether near-complete human proteome coverage could be achieved in even shorter analytical timeframes using the Orbitrap Astral Zoom MS, we fractionated a HEK293 peptide digest by offline HpH into 24, 34 and 46 fractions. Each fractionation scheme was analyzed using 180 or 300 SPD method (Fig. 5A). We compared the proteome coverage achieved for each fractionation scheme as a function of MS measurement time to 1-hour single-shot analysis (24 SPD) and the 34 fractions data published by Guzman et al. (Fig. 5B). Using a 1 h single-shot gradient, we achieved a proteome coverage of 10,247 proteins – about ∼2000 proteins short of a complete proteome. In contrast, by analyzing 34 fractions with 300 SPD multi-shot approach, the Orbitrap Astral Zoom MS significantly expanded the proteome coverage, accounting for just 2.7 h of total LC-MS analysis time. This enables the identification of > 12,100 protein groups, with the majority of those uniquely detected by the multi-shot method corresponding to low-abundance species, including membrane proteins and transcription factors (Supplementary Fig. 4A). This represents a twofold increase in throughput for comprehensive proteome profiling compared to the Orbitrap Astral MS and a fourteenfold increase compared to the Q Exactive HF-X^26^. Although we covered a comparable number of proteins with the faster 300 SPD analysis, the >200,000 unique peptides identified were ∼10% lower than with the 180 SPD method, likely reflecting the reduced chromatographic resolution inherent to shorter gradients (Fig. 5B). We achieved high overall sequence coverage, which enabled analysis of post-translational modifications (PTMs) without specific enrichment. For instance, searching the 34 fractions for site-specific phosphorylation identified 2089 phosphorylation sites adhering to the expected target amino acids with ∼85% serine, ∼15% threonine, and <1% tyrosine (Fig. 5C). The analysis of these sites using sequence logo motifs and molecular function GO term enrichment revealed a strong enrichment of substrates for proline-directed kinases, such as cyclin-dependent kinases (CDKs), which play critical roles in regulating cell cycle progression and modulating processes related to DNA replication and checkpoint control^27,28^ (Supplementary Fig. 5 B,C) In addition, the near-complete HEK293 proteome data also allowed us to identify protein variants, including alternative splicing events, single amino acid variants (SAVs), and splicing-associated variants including insertions and deletions (indels) by searching the data against a large protein database that was translated from transcriptomic information about HEK293 variants. In total, we identified 935 alternative splicing events, 743 SAVs, and 11 indels (Fig. D). Gene Ontology (GO) analysis of the detected splicing variants revealed enrichment in biological processes related to DNA repair and embryonic development, consistent with regulatory functions during cell proliferation and genome maintenance. These findings reflect the central role of alternative splicing as a key mechanism for expanding proteomic diversity, particularly within genes involved in regulatory, signaling, and developmental pathways^29^. By contrast, SAVs were predominantly enriched in mitochondrial and metabolic pathways, including the carnitine shuttle, mitochondrial electron transport chain and ciliary body morphogenesis, which can reflect genetic variation or adaptation in long-term cultured cells, particularly impacting energy-producing organelles such as mitochondria^30^ (Supplementary Fig. 5 D,E).

**Fig. 5.**
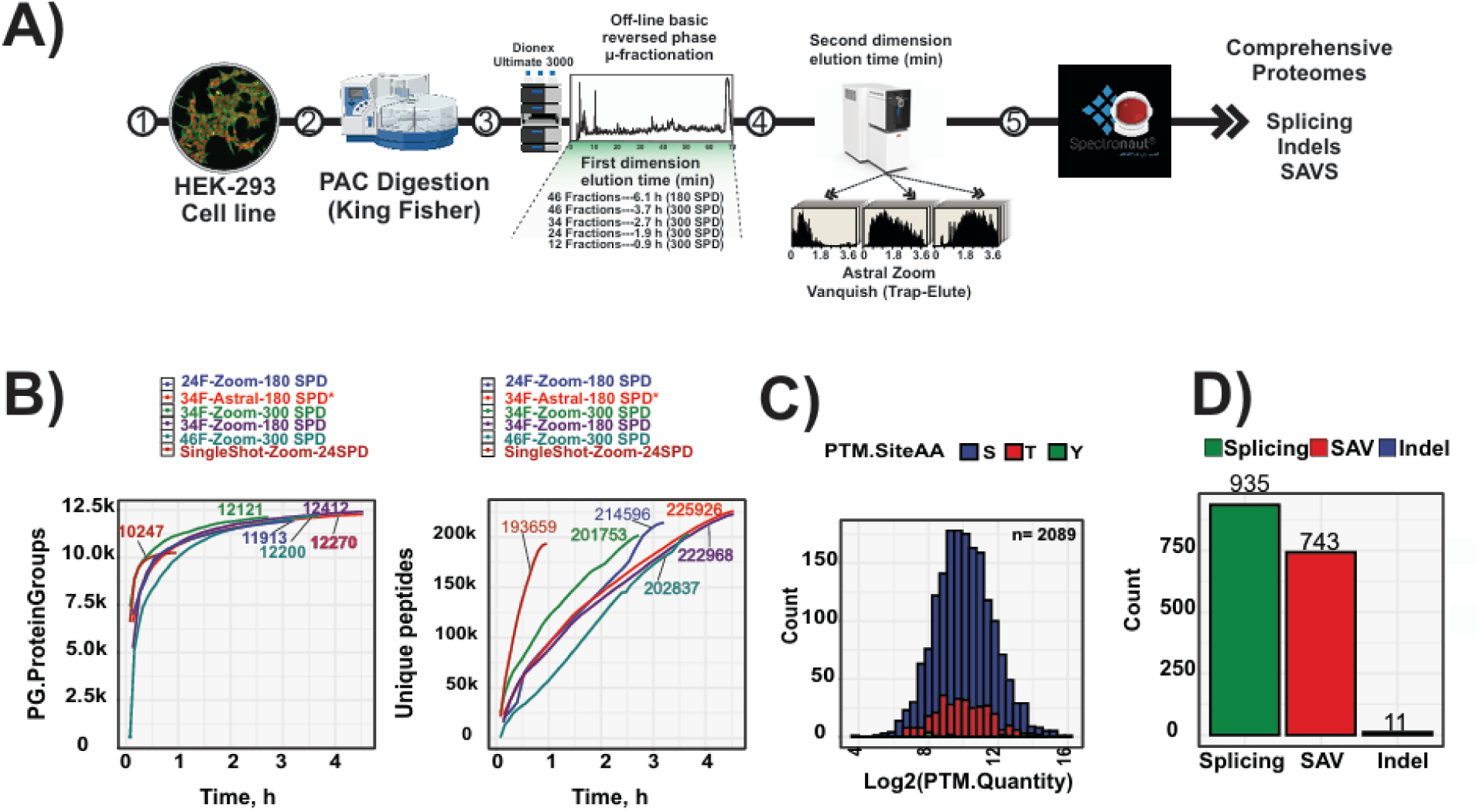
High-Coverage Human Proteome Profiling via Optimized Multi-Shot Fractionation and Fast LC-MS Workflows. **A)** Schematic of the experimental workflow illustrating the fractionation strategies optimized for comprehensive proteome coverage. **B)** Cumulative number of protein groups and peptides identified (right and left panels, respectively) as a function of total MS acquisition time. Shown are results from 46 and 34 high-pH reversed-phase (HpH) fractions acquired at 300 SPD, 34 and 24 HpH fractions acquired at 180 SPD, and single-shot analysis using a 24 SPD gradient on the Orbitrap Astral Zoom MS. For comparison, 34 HpH fractions acquired at 180 SPD using the Orbitrap Astral MS are also included^17^ (Guzman et al., 2024). **C)** Sequence logo and abundance distribution of phosphorylation sites identified without enrichment. **D)** Bar chart showing the number of detected splicing events, single amino acid variants (SAVs), and insertions/deletions (indels).

### Fast and deep proteome profiling with Orbitrap Astral Zoom MS and Short-Gradient LC

To fully leverage the acquisition speed of 270 Hz in nDIA mode, we hypothesized that the Orbitrap Astral Zoom MS would be ideally suited for use in combination with very-short LC gradients. To test this, we used a trap-and-elute LC method allowing 300 SPD analysis. Analyzing 200 ng of HEK293 tryptic digest with the fastest nDIA method of 2 Th isolation windows and 1.5 ms maximum IT, we identified ∼100,000 unique peptides and 8,400 protein groups (Fig. 6A), with excellent quantitative reproducibility across technical replicates (Pearson correlation coefficient of 0.99 for >8,400 proteins; Fig. 6B). To evaluate whether even higher-throughput proteome analysis is feasible, we employed the Evosep One LC equipped with the Eno module, which provides enhanced synchronization of valve switching and pump control. This improvement enables robust chromatographic performance, particularly under ultra-fast gradient conditions such as the 500 SPD method, making it suitable for large-scale proteomic studies involving thousands of samples, including clinical research and systems biology applications. The 500 SPD method is designed for ultra-high throughput using a 2.2 min gradient with a total run-to-run time of only 2.88 minutes. Analyzing 200 ng of HEK293 tryptic digest with the 500 SPD method, we found that the Orbitrap Astral Zoom MS identified on average 7184 unique protein groups from ∼67,000 unique peptides, which is >15,000 more peptides than covered by the Orbitrap Astral MS (Fig. 6C). This was achieved with very high reproducibility between replicates with Pearson correlation coefficient of >0.99 consistently quantifying >7000 MaxLFQ normalized proteins (Fig.6D). The 500 SPD method provides 72 seconds of active peptide elution and during the main part of this, >1,000 unique peptides per second were identified when using the Orbitrap Astral Zoom MS (Supplementary Fig. 6A). This is better illustrated by the cumulative number of unique peptides identified from the onset of the active gradient, with a linear increase in identifications and 50,000 unique peptides identified within the first 50 seconds (Supplementary Fig. 6B). As the number of peptides per protein identified is biased towards proteins of high abundance, we did not expect to see a similar linear increase in unique proteins across the gradient. Indeed, the cumulative number of unique proteins initially grew very fast with 2,000 proteins identified in 7 seconds, 4,000 proteins identified in 14 seconds, and 6,000 proteins identified in 30 seconds of active gradient, ultimately reaching saturation with ∼7,000 proteins in approximately one minute (Supplementary Fig. 6C).

**Fig. 6.**
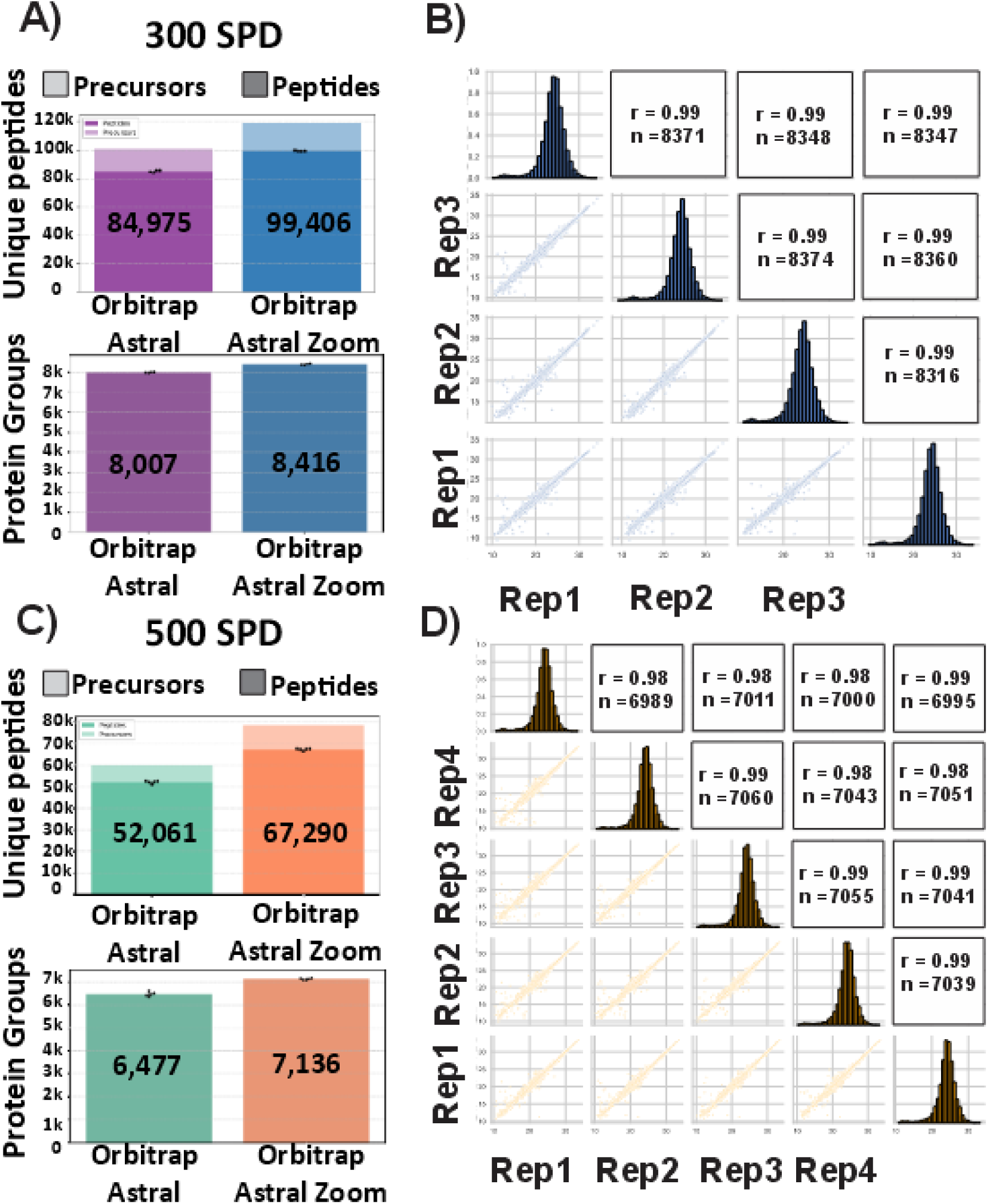
Ultra-fast and deep proteome profiling with Orbitrap Astral Zoom MS and short-gradient LC. **A)** Number of unique peptides, precursors (top), and protein groups (bottom) identified from 200 ng HEK293 digest analyzed with a 300 SPD trap-and-elute LC gradient using the Orbitrap Astral Zoom MS. **B)** Quantitative reproducibility of protein intensities across three technical replicates at 300 SPD. Shown are pairwise Pearson correlation coefficients (upper triangle), log₂ intensity histograms (diagonal), and scatterplots of protein intensities between replicate pairs (lower triangle). **C)** Same as in panel A but using a 500 SPD method on the Evosep Eno LC system coupled to the Orbitrap Astral Zoom MS**. D)** Same as in panel B, showing reproducibility across four technical replicates at 500 SPD.

### A Comprehensive Draft of the Human Proteome Atlas via Ultra-Fast LC-MS

To assess coverage and quantitative performance of fast LC gradients for human proteome profiling, we analyzed a panel of 32 human cell lines representing cell types of the major organs. All samples were analyzed with both 180 SPD and 300 SPD in quadruplicates with the Orbitrap Astral Zoom MS, and collectively we identified >11,500 proteins with high overall reproducibility between the two LC gradients (Fig.7A). Surprisingly, we found that the average protein coverage per cell line was comparable between the 180 SPD and 300 SPD with an average 9645 proteins and 9549 proteins identified, respectively (Fig. 7B). The 180 SPD LC gradient provides ∼5 minutes of active peptide elution, whereas the 300 SPD method comparatively provides about half that duration with ∼2.5 minutes. Nonetheless, we find very high correlation between cell line proteomes representing the same cell type between the 180 SPD and 300 SPD (Fig. 7C). Moreover, principal component analysis (PCA) of the cell line proteomes acquired with 180 SPD and 300 SPD gradients were almost perfectly overlayed when visualizing the two major principal components, indicating high concordance in global proteomic profiles across the two acquisition methods. (Fig. 7D). In addition, the Euclidean distance between cell lines was preserved across gradients, indicating consistent relative proteomic relationships within each acquisition method. The Gene Ontology analysis of different clusters derived from hierarchical heat map clustering confirmed consistent enrichment of biological processes across both 180 SPD and 300 SPD workflows, highlighting the consistency of the proteomic signal across gradient lengths. (Supplementary Fig. 7)

**Fig. 7.**
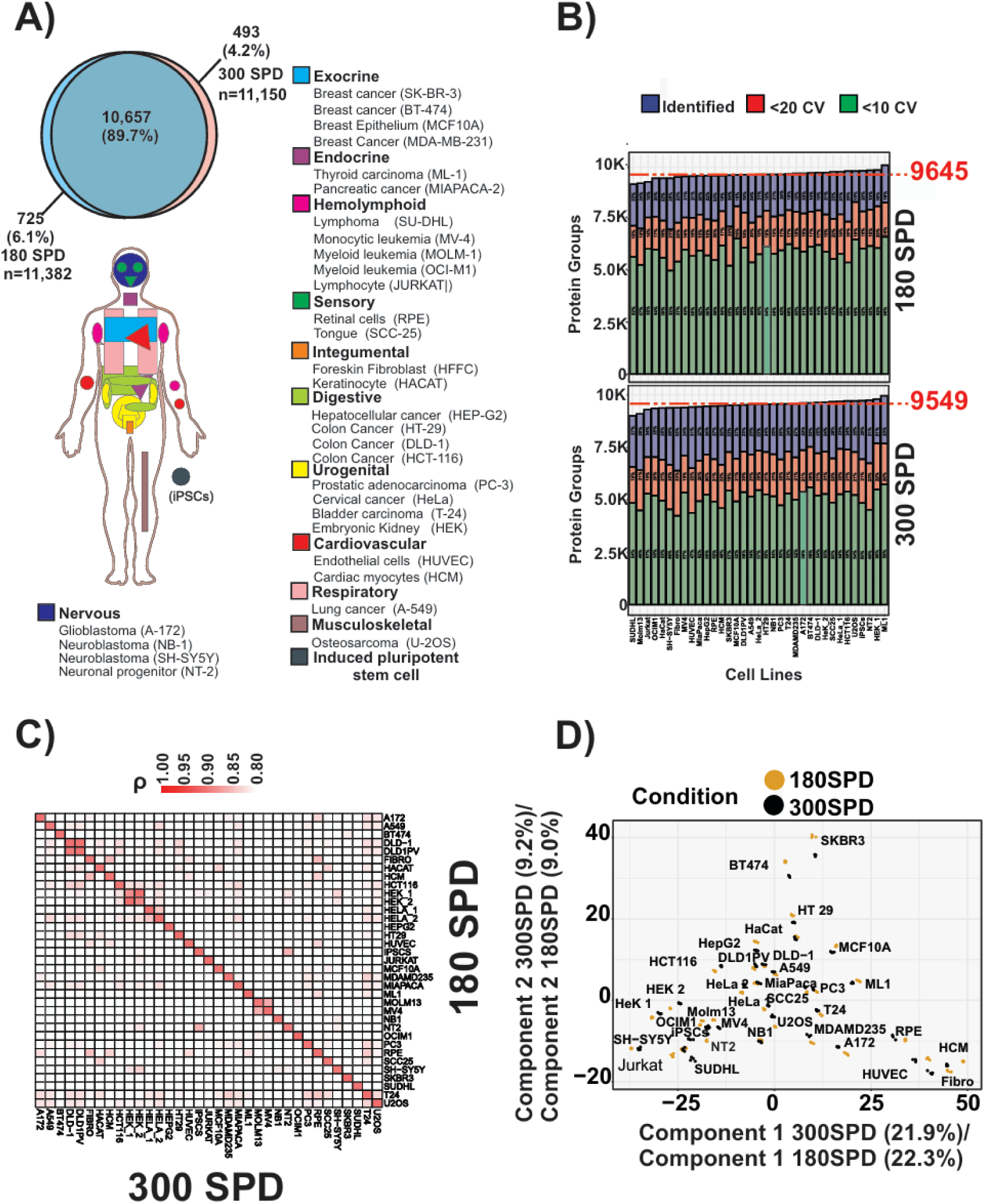
Fast and Deep Proteome Profiling with the Orbitrap Astral Zoom MS Using Short-Gradient LC. **A)** Overview of the tissue origins of 32 human cell lines profiled in technical quadruplicates using DIA with 300 SPD and 180 SPD LC-MS/MS workflows (bottom). Venn diagram (top) shows the overlap and total number of protein groups identified across both gradient conditions **B)** Bar plot of the protein groups detected at 300 SPD (right) and 180 SPD (left). Median identifications are indicated by a red dashed line. Coefficient of Variation (CV) below 10% and 20% CV (n = 4) are depicted in green and red, respectively. **C)** Pearson correlation heatmap of the 180 SPD and 300 SPD datasets. **D)** PCA analysis of the 32 cell lines recorded at 180 SPD and 300 SPD.

## Discussion

The development of the Orbitrap Astral Zoom mass spectrometer represents a significant advancement in proteomics instrumentation. By achieving ultra-fast MS/MS scan rates while maintaining high sensitivity, quantitative accuracy, and proteome depth, this mass spectrometer overcomes longstanding limitations in scalability and speed. In combination with very fast online LC gradients, this platform has the potential to redefine the landscape of proteomics research, enabling more comprehensive and efficient exploration of biological systems at an unprecedented throughput and scale. Building upon the Orbitrap Astral MS technology, the Orbitrap Astral Zoom MS introduces key advancements in duty cycle efficiency, ion-accumulation, ion-transfer, and spectral processing. These modifications result in a 30% increase in MS/MS acquisition speed, rising to 270 Hz compared to the ∼200 Hz of the Orbitrap Astral MS. The increased scan rate is accompanied by sufficient sensitivity and quantitative reproducibility to bridge the gap between data-dependent acquisition (DDA) and data-independent acquisition (DIA) methods. These advancements extend previously established Orbitrap Astral MS capabilities, by enabling faster analysis times while maintaining, or even improving, proteome coverage and quantitative reproducibility. The Orbitrap Astral Zoom mass spectrometer allows for near-complete proteome coverage within a fraction of the acquisition time required by earlier instruments. This is exemplified by the identification of >12,000 proteins in only 2.7 hours via offline high-pH reversed-phase peptide chromatography combined with very fast online 300 SPD LC-MS/MS analysis. Furthermore, the ability to analyze up to 500 samples per day with high reproducibility (Pearson correlation coefficients >0.99) opens doors for large-scale biomarker discovery, systems biology, and high-throughput drug screening. This substantial reduction in acquisition time and cost makes comprehensive proteomics more accessible, expanding its adoption makes comprehensive proteomics more accessible across diverse research fields. By dramatically increasing throughput without sacrificing data quality, the Orbitrap Astral Zoom MS enables large-scale, high-resolution proteomics studies that were previously impractical or prohibitively expensive. Applications in longitudinal studies, multi-omics integration, and single-cell proteomics are now more feasible. Additionally, the improved precision and reproducibility of the instrument enhance the reliability of datasets, making them more suitable for advanced statistical modeling and machine learning approaches, which often require high-quality, large-scale data. The Orbitrap Astral Zoom MS exemplifies the trajectory of proteomics toward increased speed and scalability without compromising sensitivity or data quality. As proteomics becomes an essential tool in systems biology, precision medicine, and clinical research, the demand for ultra-fast and reproducible workflows will continue to grow. The Orbitrap Astral Zoom MS sets the precedent for future developments, such as integrating proteomics with other omics platforms and advancing real-time proteome monitoring in clinical settings.

The technological innovations in the Orbitrap Astral Zoom MS, in particular the parallel ion pre-accumulation, faster quadrupole ion transfer, and enhanced spectral processing algorithms are central to its performance. These features complement each other, with pre-accumulation and faster ion transfer adding 66% effective injection time, facilitating an extra 70 scans per second at the same sensitivity. Furthermore, the efficacy of enhanced spectral processing enables resolving and interpretation of the more complex spectra generated when analyzing highly complex samples with slower and more sensitive methodology. Collectively, these features ensure increased performance regardless of acquisition method, enabling the instrument to achieve both increased scan rates and greater proteome depth while maintaining high sensitivity. The introduction of enhanced dynamic range (eDR) mode further expands its utility, enabling deeper peptide and protein identification with data-dependent acquisition. This technological leap ensures that the Orbitrap Astral Zoom MS is well-suited for both exploratory and targeted proteomics applications.

The Orbitrap Astral Zoom MS has already demonstrated the capacity to facilitate new research avenues. For instance, it was used to analyze 300 or 500 samples per day with high reproducibility, enabling rapid profiling of human cell lines and clinical research samples. This capability is critical for studying heterogeneous biological systems, such as cancer tissues or immune responses, where large sample sizes are required to capture variability. Moreover, the ability to perform ultra-high-throughput analyses supports the development of personalized medicine approaches by enabling the rapid identification of disease-specific biomarkers. While the instrument represents a significant advancement, certain limitations remain. For example, the ultra-short LC gradients (e.g., 300 and 500 SPD) may compromise the detection of low-abundance proteins such as cell type-specific transcription factors in highly complex samples and, although partially compensated for by higher scan rates, very short chromatographic gradients inherently limit precursor peak width and can thus inhibit precise quantification. Additionally, while the instrument achieves remarkable scan rates, the effective utilization of this capability depends on downstream data analysis pipelines that can handle the increased data volume.

Several questions remain unanswered, such as how the Orbitrap Astral Zoom MS performs in more diverse sample types, beyond standard cell lines and model organisms. Furthermore, its ability to resolve post-translational modifications (PTMs) with ultra-fast scanning methods remains to be fully explored. Finally, while the instrument has shown excellent reproducibility, its performance in longitudinal studies with highly dynamic proteomes warrants further investigation.

In summary, the Orbitrap Astral Zoom mass spectrometer represents a significant landmark advancement in proteomics technology, combining high speed, sensitivity, and scalability. It sets a new standard for large-scale biological and clinical research, with the potential to profoundly impact on medical research, particularly in the areas of biomarker discovery, disease stratification, and therapeutic monitoring. The ability to analyze large clinical research cohorts rapidly and with high reproducibility will facilitate the identification of disease-associated proteomic signatures, paving the way for real-time monitoring of patient proteomes. In the long term, this technology could support the development of personalized medicine approaches by providing detailed, patient-specific proteomic insights. While challenges remain, the instrument sets the stage for the next generation of proteomics research, enabling unprecedented insights into the complexity of biological systems.

## Materials and Methods

### Sample preparation

#### Cell lines

All human cell lines were cultured in DMEM (Gibco, Invitrogen), supplemented with 10% FBS, 100 U ml−1 penicillin (Invitrogen), and 100-μg ml−1 streptomycin (Invitrogen). The cell cultures were maintained at 37 °C, in a humidified incubator with 5% CO_2_. Cells were collected at approximately 70% confluence by washing twice with PBS (Gibco, Life technologies). Subsequently, boiling lysis buffer (5% SDS, 5 mM Tris(2-carboxyethyl)phosphine, 10 mM chloroacetamide, 100 mM Tris, pH 8.5) was added directly to the plates. The cell lysate was collected by scraping the plates and boiled for an additional 10 min. Following this, micro-tip probe sonication (Vibra-Cell VCX130, Sonics) was performed for 2 min with pulses of 1 s on and 1 s off at 80% amplitude. Protein concentration was determined by BCA method.

#### Preparation of samples for LC–MS/MS analysis

The human cell lines were digested overnight at 37 °C with looping mixing, employing the Protein Aggregation Capture protocol^31^ with MagReSyn® amine microparticles (ReSyn Biosciences) on a fully automated KingFisher platform. Briefly, 1 mg of protein lysate was resuspended in acetonitrile (ACN) to a final concentration of 70%. MagReSyn amine microparticles were added in a proportion 1:2 (protein:beads). The proteolytic digestion was performed by addition of lysyl endopeptidase (LysC, Wako) and trypsin enzymes at 1:500 and 1:250 protein ratio, respectively, to 300 µl of digestion buffer (50 mM Ammonium Bicarbonate). Protein aggregation was carried out in two cycles, each consisting of 1 min mixing at 1000 rpm followed by a 10-min pause. The beads were subsequently washed three times with 1 ml 95% ACN and two times 1 ml 70% ACN. The resulting peptide mixture was acidified with trifluoroacetic acid (TFA) to a final concentration of 1% to quench the protease activity. Finally, the peptides were concentrated by a Sep-Pak C18 96-well Plate (Waters, Milford, MA), with peptides eluted sequentially using 150 µl of 40% acetonitrile followed by 150 µl of 60% acetonitrile and collected in a 96-weel plate. The combined eluate was dried down via SpeedVac vacuum concentrator (Eppendorf, Germany). Finally, the peptide mixture was resuspended 100 uL in 0.1 % TFA and the final peptide concentration was determined by measuring absorbance at 280 nm on a Thermo Scientific NanoDrop 2000C spectrophotometer. The protein digests from the mixed species for the LFQ analysis were purchased from Pierce for HeLa (88328), Promega for yeast (V7461) and Waters for E. coli (SKU: 186003196). They were mixed manually in three different ratios, E05-H50-Y45, E45-H50-Y05 and E25-H50-Y25, respectively. Samples were kept at −20 °C until further use.

#### Off-line HpH reversed-phase HPLC fractionation

HEK293 peptides (200 µg) were separated by HpH reversed-phase chromatography using a reversed-phase Acquity CSH C_18_ 1.7 μm × 1 mm × 150 mm column (Waters) on a Thermo Scientific UltiMate 3000 HPLC system with the Chromeleon software. The instrument was operated at a flow rate of 30 μl min^−1^ with buffer A (5 mM ABC) and buffer B (100% ACN). Peptides were separated by a multi-step gradient as follows: 0–10 min 6.5% B to 15% B, 10–59.5 min 15% B to 30% B, 59.5–67 min 30% B to 65% B, 67–70 min 65% B to 80% B, 70–77 min 80% B, 78–87 min 6.5% B. A total of 46, 24 and 12 fractions were collected at 60-s intervals. Samples were acidified using 30 µl of 10% formic acid. The samples were dried down using a SpeedVac vacuum concentrator. Sample concatenation was performed manually. 200 ng of each fraction was injected for LC–MS/MS analysis.

#### LC–MS/MS analysis

LC–MS/MS analysis was performed on an Orbitrap Astral Zoom MS prototype coupled to a Thermo Scientific Vanquish Neo UHPLC or an Evosep ENO system and interfaced online using an EASY-Spray/Nano-Flex ion source. Depending on the gradient utilized, different setups were employed, including either trap-and-elute or direct injection into commercial or home-packed analytical columns. The choice of column type was made according to the gradient employed (Supplementary Table X). A blank run of equal gradient length was run after three or four runs.

For the Data-Dependent Acquisition (DDA) experiments, the Orbitrap Astral Zoom MS prototype was operated with a fixed cycle time of 0.6 s and with a full scan range of 400– 900 *m*/*z* at a resolution of 240,000. The automatic gain control (AGC) target value for the Orbitrap analyser was set to 5e6. Precursor ion selection width was kept at 2 Th and peptide fragmentation was achieved by HCD (Normalized Collision Energy (NCE) 30%). Fragment ion scans were recorded at a resolution of 80,000 @ m/z 524, with a maximum fill time (maxIT) of 3 ms. Dynamic exclusion was enabled and set to 10 seconds. The AGC target value for the Astral analyzer was set to 2e4. In Wide Window Data-Dependent Acquisition (WWDDA) experiments, the same settings were applied, except the precursor ion selection width was maintained at 4 Th, and fragment ion scans were recorded with a maxIT of 6 ms. When the enhanced dynamic range mode was used, five symmetric MS1-windows of 100 Th each were configured, covering the range of 400-900 *m/z*, with an AGC target value of 5e6 and a maxIT of 50 ms on the Orbitrap analyzer. The precursor ion selection width was set to either 2 Th or 4 Th, and MS/MS spectra were recorded with maxIT values of 3 ms and 6 ms, respectively.

For the Data-Independent Acquisition experiments, an Orbitrap Astral Zoom MS prototype was operated at a full-MS resolution of 240,000 with a full scan range of 380– 980 *m/z* otherwise stated. The full-MS AGC target value was set to 5e6. Fragment ion scans were recorded at a resolution of 80,000@m/z524 and maxIT of 1.5 ms. We used 300 windows of 2 Th scanning from 380 to 980 *m*/*z*, unless stated otherwise in Supplementary Table X. The isolated ions were fragmented using HCD with 25% NCE. For the HEK293 measured at 500 SPD in the Orbitrap Astral mass spectrometer, peptides were eluted online from the EvoTip using an Evosep ENO system (Evosep Biosystems) using a commercial - cm analytical column (EVOSEO (Endurance EV-1107 4cm x 150 um, 1.9 um). The mass spectrometer was operated in positive mode using the DIA mode. Full scan precursor spectra (380–980 *m/z*) were recorded in profile mode using a resolution of 240,000 at *m*/*z* 200, with an AGC target value of 5e6 and a maxIT of 5 ms. Fragment spectra were then recorded in profile mode, fragmenting 300 consecutive 2 Th windows covering the mass range 380–980 *m/z*. Isolated precursors were fragmented in the HCD cell using 25% NCE, and an AGC target value of 2e4..

Further details are described in Supplementary Table X.

#### Raw MS data analysis

Raw files from DIA and DDA comparison experiments were analyzed in DIA-NN 1.9.2^32^ software allowing for C carbamidomethylation and N-terminal M excision and 1 missed cleavage). The spectral library was generated from a human reference database (UniProt 2024 release, 20,417 sequences). Raw files from single-shot dilution series of HEK peptides, optimization and cell lines were analyzed in Spectronaut v.19 (Biognosys) with a library-free approach (directDIA+) using the human reference database (UniProt 2024 release, 20,417 sequences) complemented with common contaminants (246 sequences). Cysteine carbamylation was set as a fixed modification, whereas methionine oxidation and protein N-terminal acetylation were set as variable modifications. Precursor filtering was set as *Q* value, and cross-run normalization was unchecked. Each experiment was analyzed separately, and those that contained different experimental conditions (different input amounts or acquisition methods) were searched, enabling method evaluation and indicating the different conditions (each one with *n* = 4 experimental replicates) in the condition setup tab.

Raw files from LFQ analysis of the mixed species samples were analyzed in Spectronaut v.19 (Biognosys) with a library-free approach (directDIA+) using a benchmark reference database for the three species (31,657 sequences in total). Cysteine carbamylation was set as a fixed modification, whereas methionine oxidation and protein N-terminal acetylation were set as variable modifications. Precursor filtering was set as *Q* value, and cross-run normalization was enabled. Each experiment consisting of samples with the same loading amounts was analyzed separately, and each condition contained three experimental replicates.

Raw files from different fractionation schemes were analyzed in Spectronaut v.19 (Biognosys) with a library-free approach (directDIA+) using the human reference database (UniProt 2024 release, 20,417 sequences) complemented with common contaminants (246 sequences). Methionine oxidation and protein N-terminal acetylation were set as variables, whereas cysteine carbamylation was set as a fixed modification. Precursor filtering was set as *Q* value. Each fractionation scheme was searched independently, except for searches performed in triplicate. Quantification was performed using the MaxLFQ algorithm embedded in iq R package^33^. Briefly, extended Spectronaut output results were filtered as follow: PG.Qvalue < 0.01 and EG.Qvalue < 0.01. Then, the MaxLFQ algorithm was applied using PG.Genes and PG.ProteinNames for protein annotation. Finally, to determine the percentages of residues in each identified protein sequence (sequence coverage), the program Protein Coverage Summarizer was used. The human reference database used for quantification and a file containing all detected peptide sequences with a protein name associated (PG.Qvalue < 0.01 and EG.Qvalue < 0.01) were utilized for protein assembly and sequence coverage calculation. For the phosphopetides search, the 34 fractions scheme was analyzed in triplicate using an empirical library generated in-house by the HpH fractionation (12 fractions) of phosphopeptide enrichment (119,793 precursors). Spectronaut output was reformatted using the Perseus plugin^34^ peptide collapse to create a MaxQuant-like site-table. MS statistical plots were based on MS1 feature detection output retrieved by MaxQuant (v.1.6.7.0) or MaxQuant (v.1.6.14.0). Representative raw files for each method were loaded into MaxQuant and analyzed without the indication of a FASTA file, to extract only the relevant MS features. Enrichment analysis was performed with the package clusterProfiler^35^ using the org.Hs.eg.db. All data analysis was performed using R v.4.2.2 and R studio v.2022.12.0 Build 353.

Dia-NN analysis was performed using 1.9.2^32^, either as Library free or with a Library. All Dia-NN output files were filtered for protein or peptide Q-value to approach 1% FDR.

For Library free analysis the standard settings were kept; pep length 7-30, Precursor Charge Range 1-4, Precursor M/Z range 300-1800, Fragment M/Z range 200-1800, 1% Precursor FDR with No shared spectra and heuristic protein inference. The Fasta file was generated from the Human proteome found on Uniprot 13^th^ of November 2024.

For the library search, a library was generated with Dia-NN from the 24, 34, 46 fraction experiment using the same FASTA file and MBR. This Library was used for Dia-NN analysis with the same settings as described in the library free method, except MBR was turned off.

Scan intensity output and injection times were extracted using inhouse scripts reading directly from the .raw file using Thermo Rawfile Reader through Python.

## Author contributions

U.H.G., I.A.H., M.R. and J.V.O. designed the proteomics experiments, U.H.G. prepared and performed proteomics experiments and U.H.G., M.R and I.A.H. analyzed the resulting data. E. Damoc. and T.N.A. helped optimize the scanning methods and evaluate the data. C.K. and O.Ø. provided samples. H.S., E.D., B.H., J.P, A.K., Y.M, I.C., J.K., K.L.F., E.C., J-P.H, D.H,, V.Z., C.H. contributed to the development of the instrument. J.V.O. designed the experiments and critically evaluated the results. U.H.G., I.A.H, M.R. and J.V.O. wrote the first draft of the paper. All authors read, edited and approved the final version of the paper.

## Inclusion and ethics statement

We are committed to promoting diversity and inclusion in science and ensuring that our research is conducted ethically and responsibly. Our study was designed and conducted in accordance with ethical principles and guidelines, including obtaining informed consent from all participants and complying with relevant regulations and laws.

## Supplementary Materials

**Supplementary Fig. 1.**
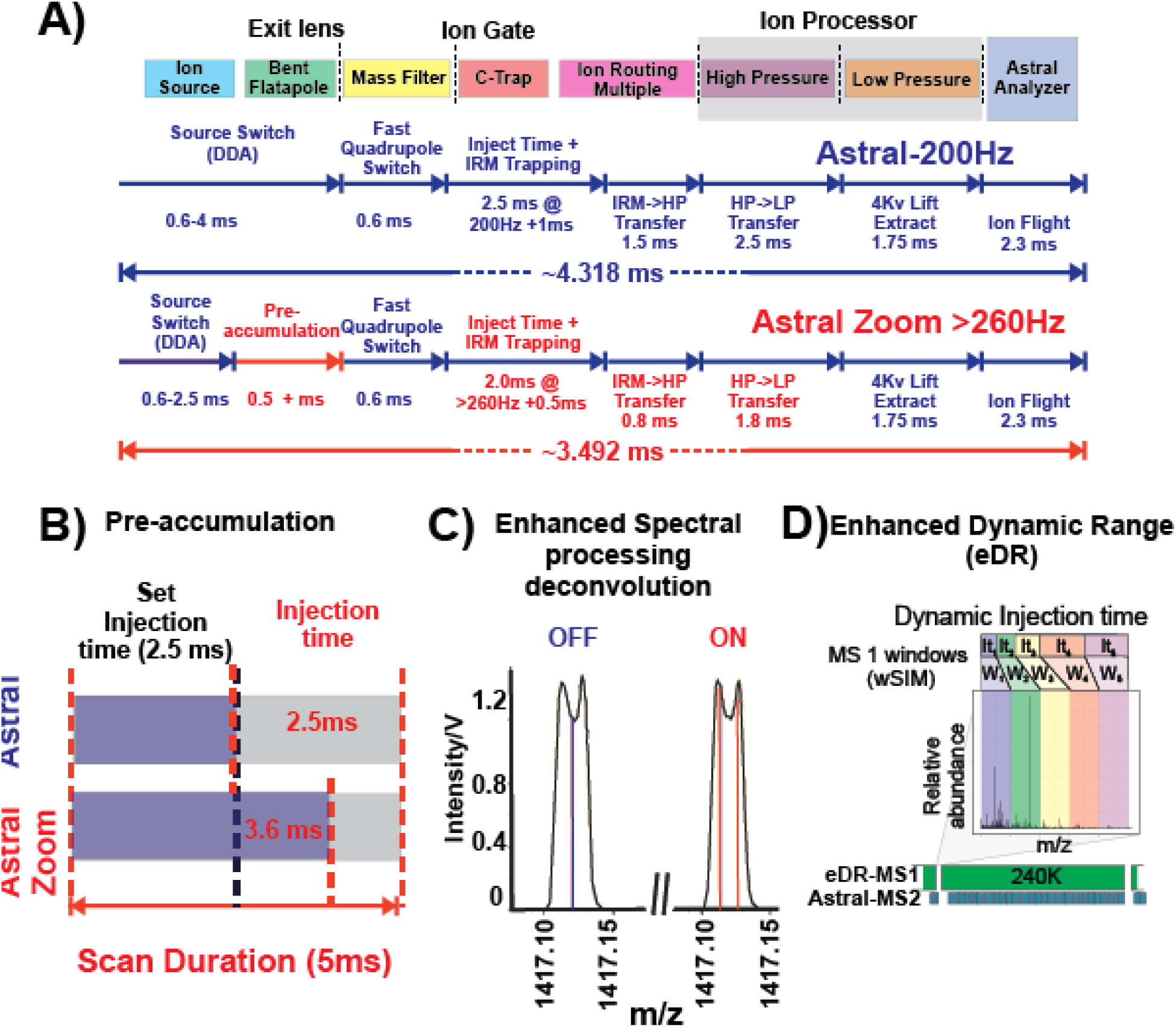
Schematics of the Orbitrap Astral Zoom Mass Spectrometer’s Ion Processing Features. **A)** Ion processing scheme depicting the improvements made to parallelized stages and timings of the Orbitrap Astral Zoom MS compared to the Orbitrap Astral MS. Median scan-scan duration is depicted for both instruments. **B)** Scheme illustrating the Pre-Accumulation feature concept when the injection time is set to 2.5 ms and no HAR enabled. **C)** Profile spectrum of a split ion peak with centroid positions when the Enhanced Spectral deconvolution algorithm (ESP) is enabled or disabled. **D)** Schematic illustrating the enhanced dynamic range (eDR) mode, featuring multiplexed ion injections with distinct accumulation times tailored to specific m/z ranges.

**Supplementary Fig. 2.**
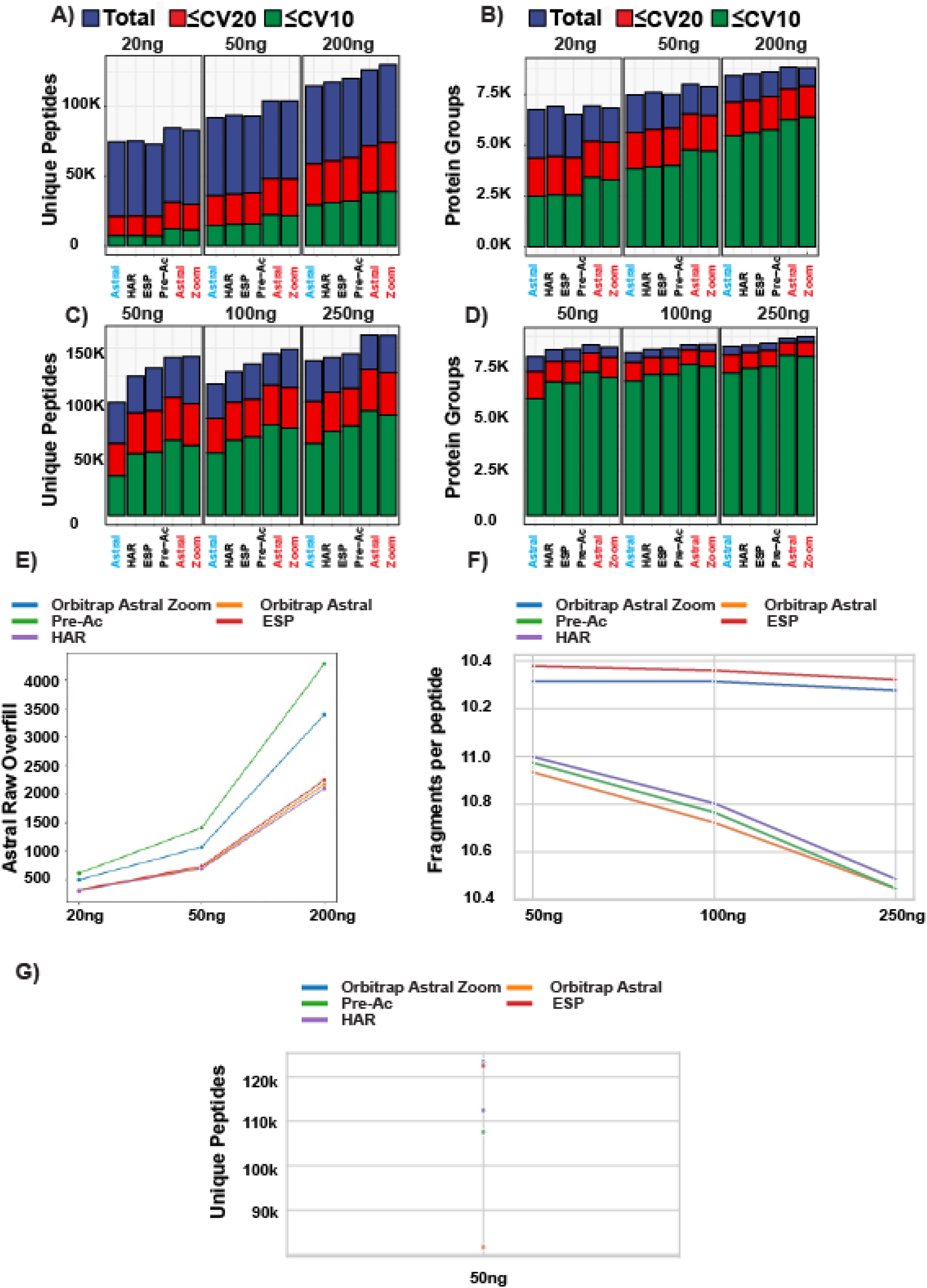
Performance Benchmarking of the Orbitrap Astral Zoom MS Features. **A,C)** Number of unique peptides and **B,D)** number of protein groups identified across a dilution series of HEK293 peptides (20–200 ng or 20–250 ng) using 90 SPD and 50 SPD gradients. The upper panels represent data with 2 Th isolation windows and 1.5 ms injection time, while the lower panels show results with 4 Th isolation windows and 6.0 ms injection time. The performance of individual features enabled on the Orbitrap Astral Zoom MS is compared to that of the Orbitrap Astral MS. Coefficients of variation (CV) below 10% and 20% across five replicates (n = 5) are indicated in green and red, respectively. Feature abbreviations: HAR, high acquisition rate; Pre-Ac, pre-accumulation; ESP, Enhanced spectral processing. **C)** Average scan-level ion overfill observed during 90 SPD acquisitions across 20-200 ng tryptic peptides from HEK293 with individual features enabled on the Orbitrap Astral Zoom MS and Orbitrap Astral MS. **D)** Average number of fragment ions per peptide identified across a dilution series (50–250 ng) of HEK293 tryptic peptides with individual features enabled during 50 SPD LC gradients (4 Th isolation windows, 6.0 ms injection time), compared to Orbitrap Astral Zoom MS and Orbitrap Astral MS. **E)** Unique identified peptides count from 50 ng HEK293 tryptic peptide injections with individual features enabled during 50 SPD LC gradients (4 Th isolation windows, 6.0 ms injection time), benchmarked against the Orbitrap Astral MS and Orbitrap Astral Zoom MS.

**Supplementary Fig. 3.**
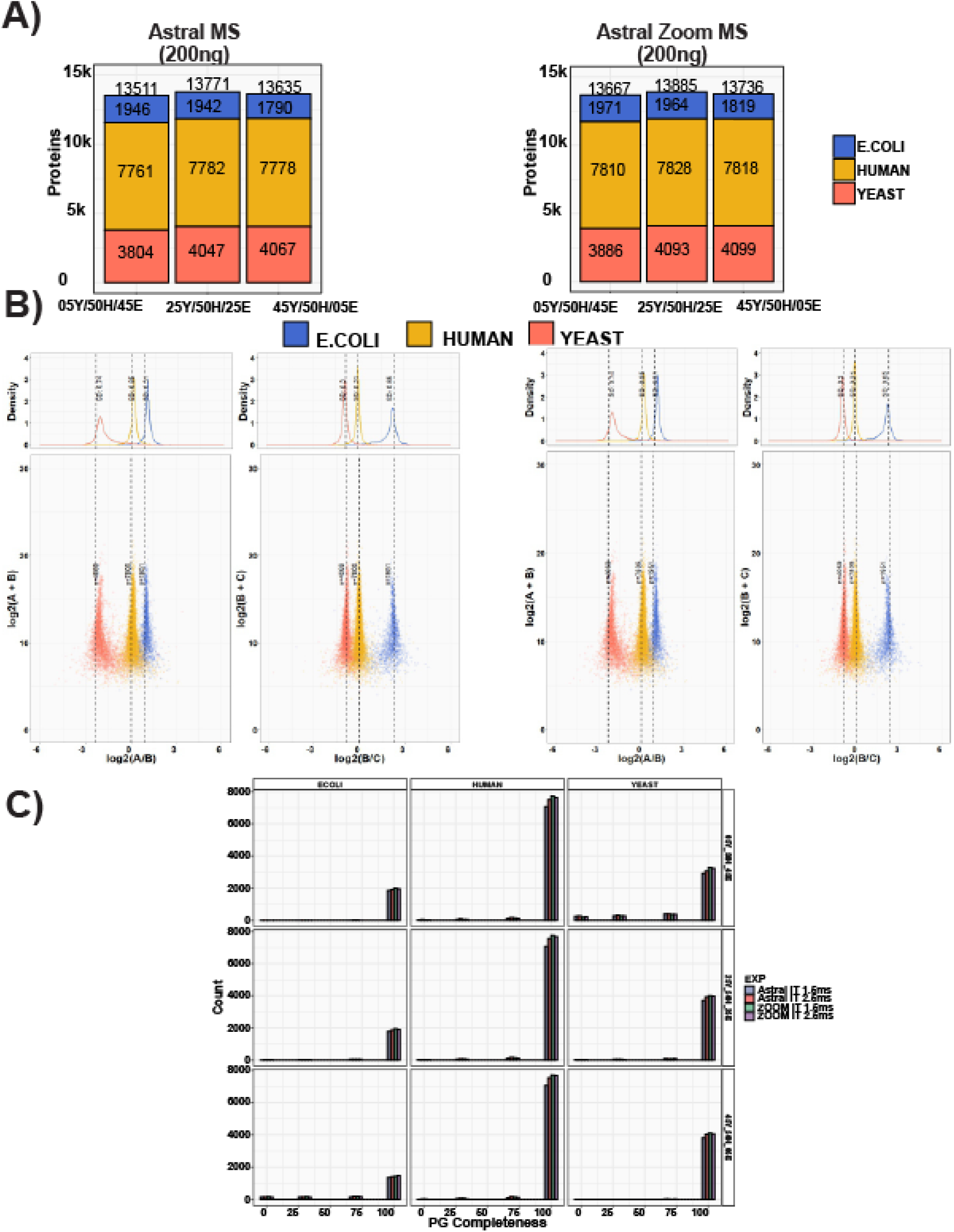
Quantitative Assessment of LFQ Accuracy and Precision on the Orbitrap Astral Zoom MS under High-Speed Scanning Regimes (2.5 ms). Samples were processed using the Orbitrap Astral MS and Orbitrap Astral Zoom MS in technical triplicates, employing a 2.5-ms maxIT and 2-Th window size method. The loading amounts were 200 ng. **A)** Number of proteins identified from the three species in each sample. For the Orbitrap Astral MS and Orbitrap Astral Zoom MS. **B)** log-transformed ratios of quantified proteins. Scatter plots for all runs over the log-transformed protein intensities are displayed at the bottom, while density plots are on the top. Colored dashed lines represent expected log2(A/B) values for proteins from humans (yellow), yeast (orange) and E. coli (blue). Standard deviations are displayed on the density plots. **C)** Data completeness is depicted for each ratio and organism.

**Supplementary Fig. 4.**
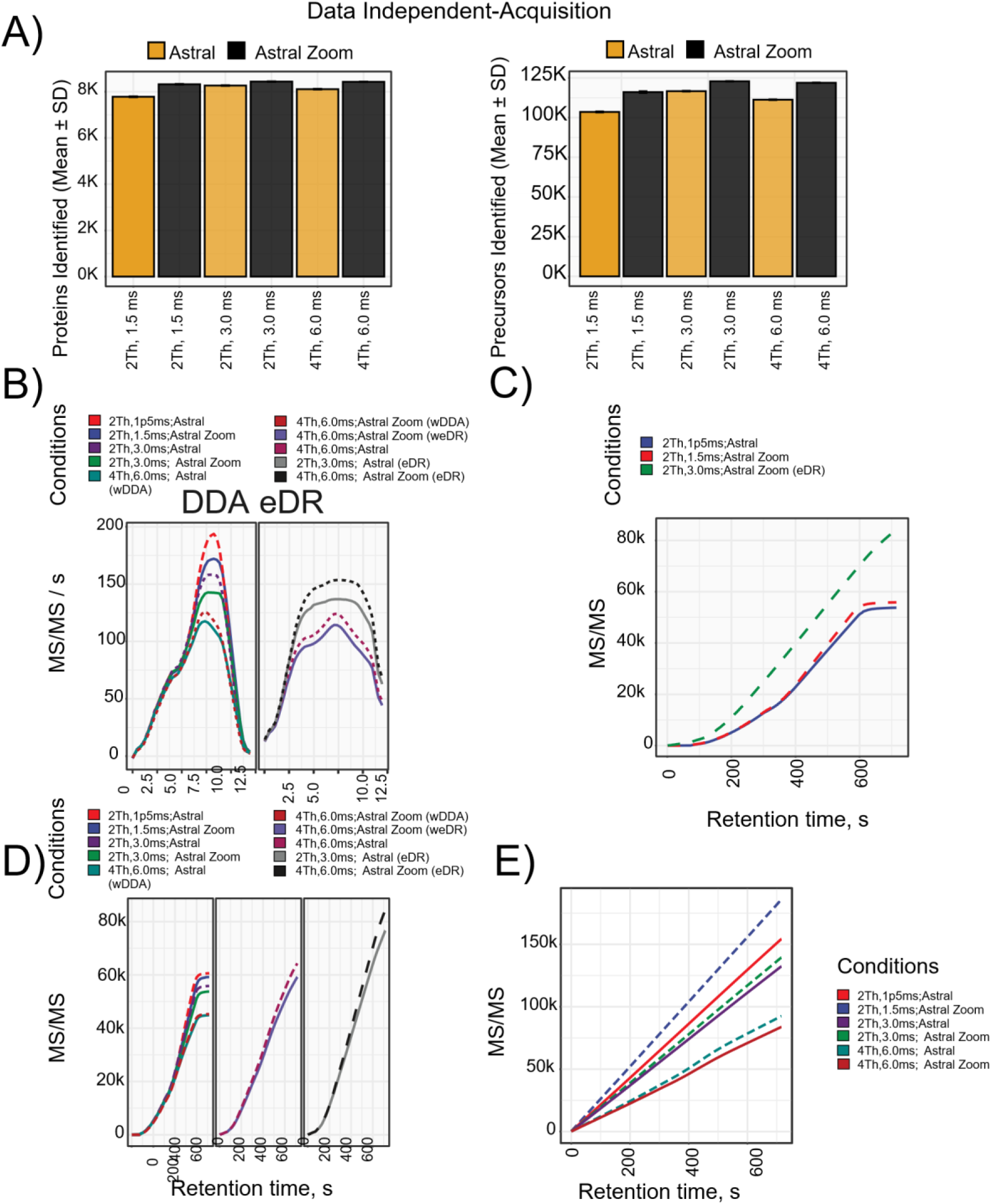
Performance benchmarking of the Orbitrap Astral Zoom MS in DDA and DIA acquisition modes. **A)** Data-Independent acquisition (DIA) performance at the protein group and peptide levels comparing the Orbitrap Astral Zoom MS to Orbitrap Astral MS. Analyses were performed at 100 SPD using 10 ng HEK 293 tryptic peptide input across four distinct DIA methods. **B)** MS/MS acquisition rate (scans per second) over the LC gradient for various DDA methods. **C-D)** Cumulative MS/MS scan counts as a function of retention time, comparing DDA methods on the Orbitrap Astral and the Orbitrap Astral Zoom MS with eDR mode on and off. **E)** Cumulative MS/MS scan counts over LC retention time for various DIA acquisition strategies, comparing fast- and high-sensitive methods across the Orbitrap Astral MS and the Orbitrap Astral Zoom MS.

**Supplementary Fig. 5.**
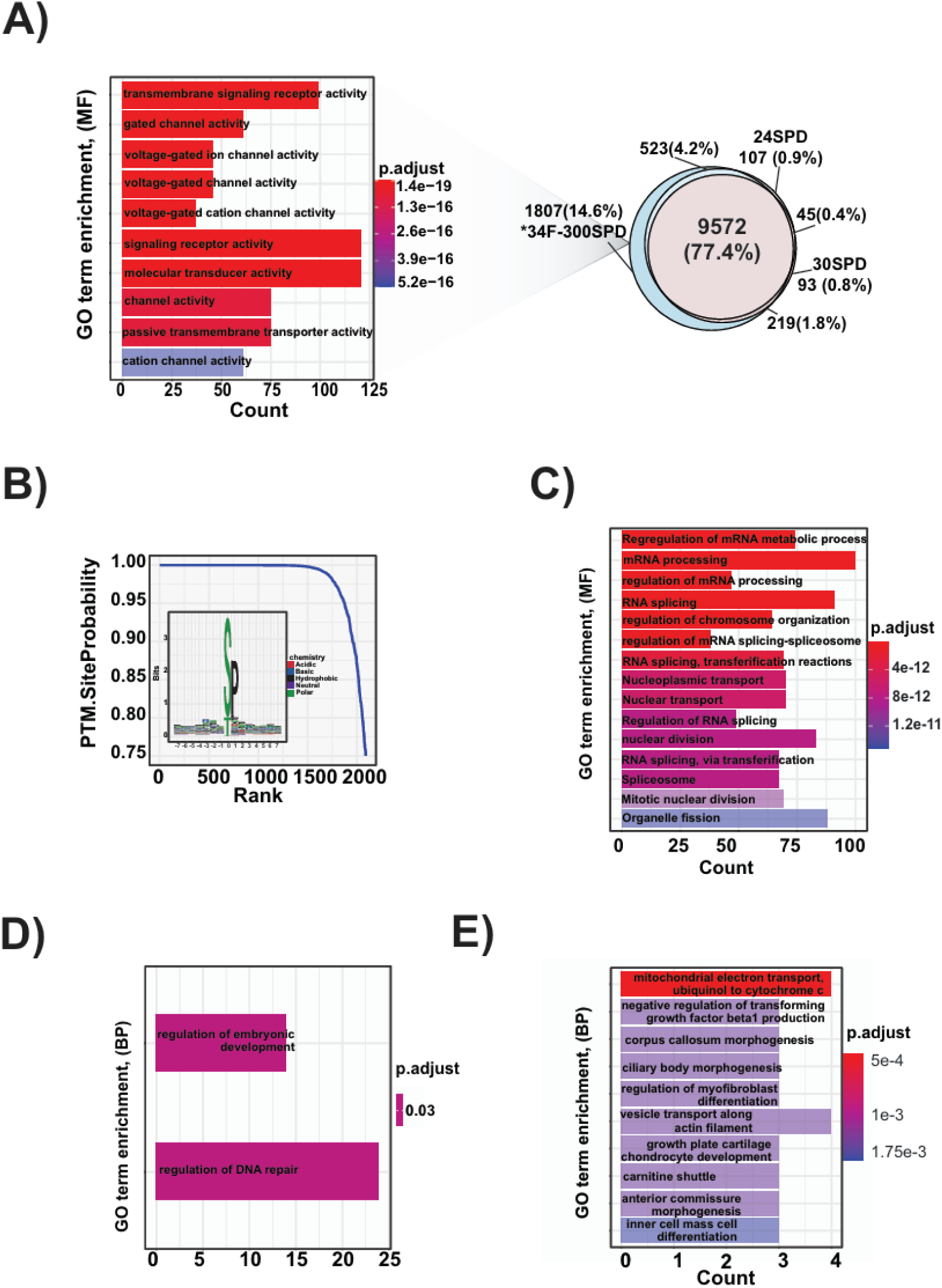
Multi-shot proteomics strategy enables detection of low-abundance peptides, post-translational modifications, and protein variants. **A)** Gene Ontology (GO) enrichment analysis of *Molecular Function* terms for proteins uniquely identified using the multi-shot approach compared to single-shot acquisition at 24 SPD. **B)** GO enrichment analysis of *Molecular Function* terms for phosphorylated proteins identified without enrichment using 34 high-pH reversed-phase (HpH) fractions acquired at 300 SPD. **C-D)** GO enrichment analysis of *Biological Process* terms associated with splice variants (**C**) and single amino acid variants (SAVs; **D**) detected using the multi-shot proteomics strategy with 34 HpH fractions.

**Supplementary Fig. 6.**
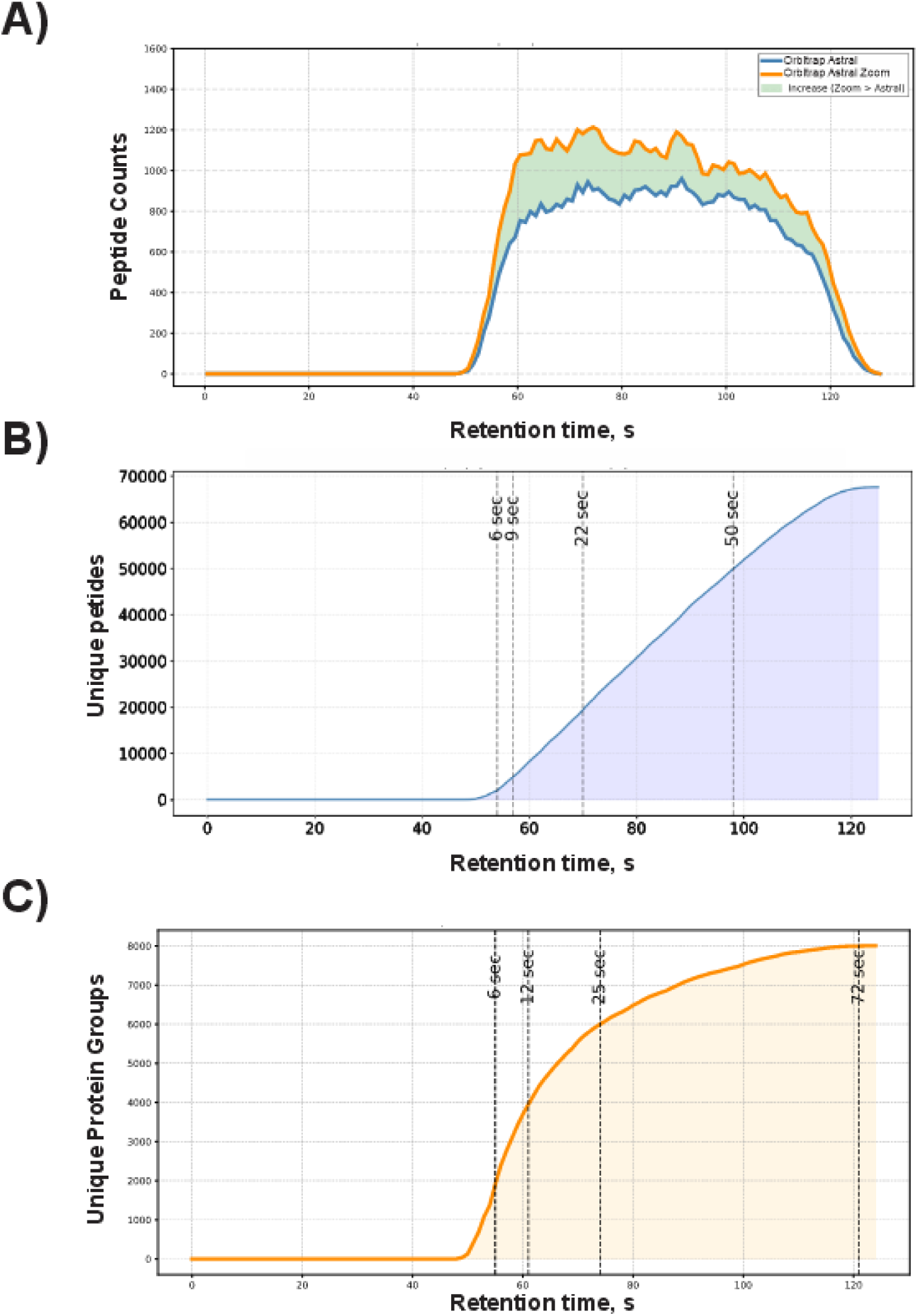
Proteome depth achieved with high-throughput ultra-fast LC gradients. **A)** Number of peptides identified over the course of the 500 SPD LC gradient for the Orbitrap Astral MS (blue) and Orbitrap Astral Zoom MS (orange); time intervals with increased identification rates in the Orbitrap Astral Zoom MS are highlighted in green. **B)** Cumulative number of unique peptides identified across the LC gradient. Dashed lines indicate the time points at which 20,000 and 50,000 unique peptides were detected. **C)** Cumulative number of protein groups identified across the LC gradient. Dashed lines indicate the time points corresponding to the identification of 2,000, 4,000, 6,000, and 8,000 unique protein groups.

**Supplementary Fig. 7.**
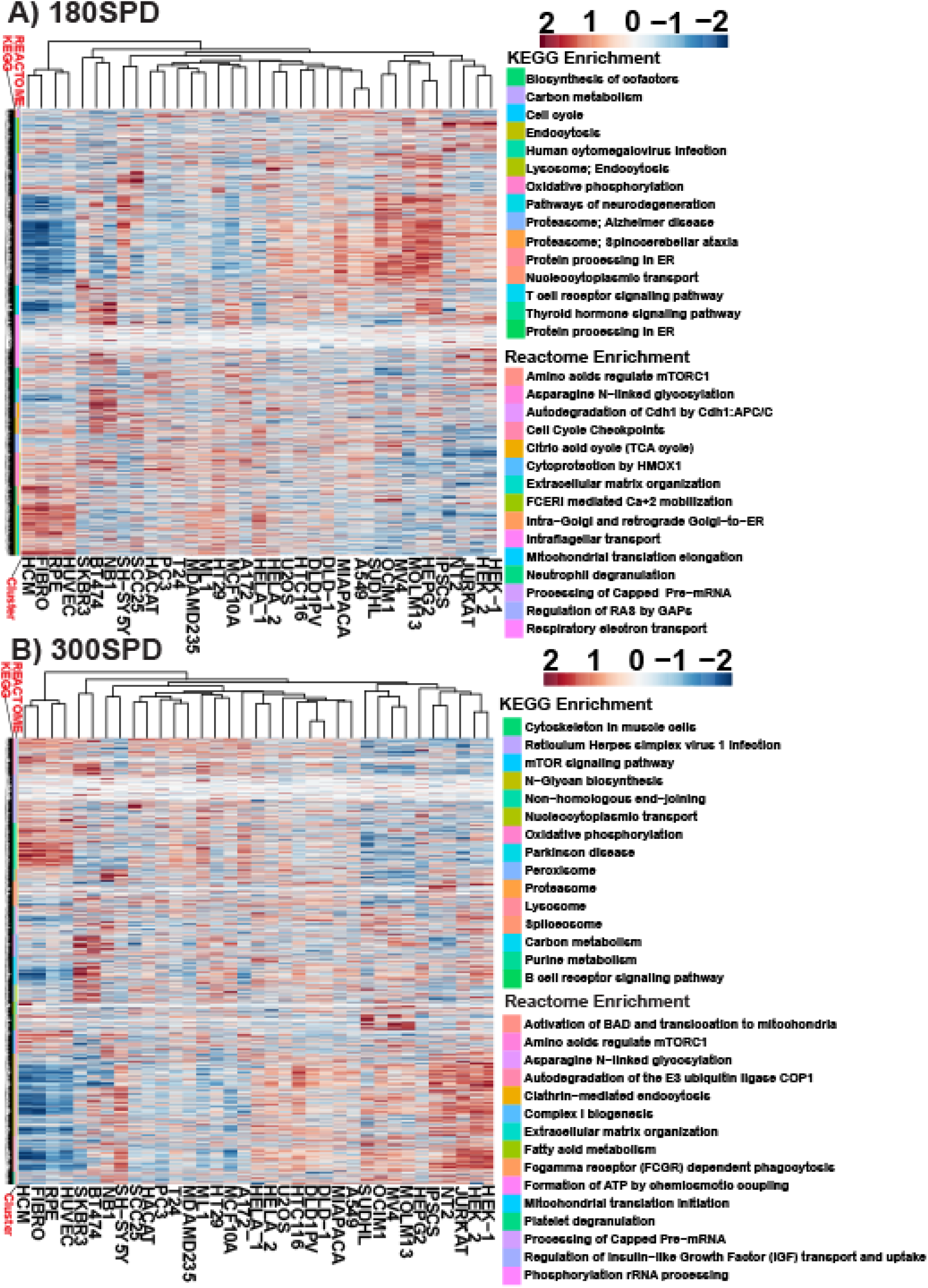
Functional landscape across 32 cell lines. **(A)** Hierarchical clustering heatmap of protein expression profiles measured across 32 cell lines at 180 SPD. Bar colors depict the KEGG and Reactome pathway enrichment analyses of the identified protein clusters associated to functional signatures. **(B)** Hierarchical clustering heatmap of protein expression profiles measured across 32 cell lines at 300 SPD. Bar colors depict the KEGG and Reactome pathway enrichment analyses of the identified protein clusters associated to functional signatures.

